# A 20-year time-series of a freshwater lake reveals seasonal dynamics and environmental drivers of viral diversity, ecology, and evolution

**DOI:** 10.1101/2024.02.06.579183

**Authors:** Zhichao Zhou, Patricia Q. Tran, Cody Martin, Robin R. Rohwer, Brett J. Baker, Katherine D. McMahon, Karthik Anantharaman

## Abstract

Long-term ecological studies are powerful tools to investigate microbiomes and ecosystem change but have mostly ignored viruses. Here, we leveraged a 20-year time-series of a freshwater lake to characterize 1.3 million viral genomes over time, seasonality, and environmental factors. We identified 578 auxiliary metabolic gene (AMG) clusters representing over 150,000 AMGs, the most abundant of which, including *psbA* for photosynthesis, *pmoC* for methane oxidation, and *katG* for hydrogen peroxide decomposition, were consistently represented in viruses across decades and seasons. We observed positive associations and niche differentiation between virus-host pairs during seasonal change including in keystone taxa, Cyanobacteria, methanotrophs, and Nanopelagicales. Environmental constraints, specifically inorganic carbon and ammonium influenced viral abundances over time, and highlighted roles of viruses in both “top-down” and “bottom-up” interactions. Key evolutionary processes shaping gene and genome-wide selection included favored fitness genes, reduced genomic heterogeneity, and dominant sub-populations carrying specific genes. Overall, our study advances understanding of diversity, ecological dynamics, and evolutionary trajectories of viruses in Earth’s microbiomes and ecosystems.

## Main

Viruses that infect bacteria and archaea (phages) are the most abundant biological entities in ecosystems. Phage can reshape microbial metabolism, drive nutrient cycling, and influence global biogeochemical cycles^1, 2^. Uncultivated viral genomes obtained from metagenomes have significantly enriched the collection of viruses in public databases and improved our understanding of viruses in nature^3^. In the largest public viral database to date (IMG/VR v4^3^), freshwater lake viruses accounted for approximately 15% of all viral genomes, ranking them first among all environmental subtypes. Despite the significant virus sequence deposition compared to other environments like oceans and soil, viruses in freshwater lakes remain understudied. Globally, freshwater lakes are undergoing rapid change due to landscape and climate alterations. Microbial communities are foundational players in freshwater ecology^4^, and characterizing the diversity, function, ecology, and evolution of their viruses will improve our understanding of a major top-down control of microbial communities.

Recently, time-series studies have been adopted in microbial ecology and have revealed microbial dynamics, population variation, and ecological impacts of microorganisms on natural ecosystems. In addition to allowing observations and analyses of temporal variation, time-series studies can also contribute to an understanding of evolutionary progression. For example, a 6-year time-series of lake pelagic bacterial community composition in Lake Mendota (Wisconsin, US) highlighted regular interannual dynamics and the link between the microbial community and seasonal drivers, which reflected the climate variation^5^. By harnessing the advantage of tracing population microdiversity changes, a 9-year time-series study of lake microorganisms in Trout Bog Lake (Wisconsin, US) revealed the evolutionary processes of bacterial speciation, which were subjected to two distinctive evolutionary models coexisting in the same environment^6^. However, few time-series studies have been conducted to date involving a comprehensive characterization of viral communities, except for a recent ecogenomic characterization of virophages^7^. By harnessing time-series metagenomics of Lake Mendota and Trout Bog Lake in Wisconsin, USA, the characterization of 25 uncultivated virophages revealed virus-host relationships between virophages and giant viruses, and unraveled ecological and evolutionary patterns over multiple years^7^.

There are known viral associations with geochemistry, primarily through the activity of auxiliary metabolic genes (AMGs)^8, 9, 10^. Functional and metabolic reprogramming of host metabolism by AMGs can maintain, drive, or short-circuit important metabolic steps, providing viruses with fitness advantages^9, 11^. The involvement of AMGs in freshwater ecosystems has been rarely reported, except for recent studies of methane oxidation and photosynthesis in freshwater lakes^12^, as compared to well-studied instances of photosynthesis^13, 14, 15^, sulfur oxidation^10, 16, 17^, ammonia oxidation^18^, and ammonification^19^ in the oceans. Additionally, research linking how viral populations and their ecological functions are influenced by environmental factors remains elusive.

Here, we leveraged time-series metagenomes collected over 20 years (2000-2019) to study freshwater viral diversity, ecology, and their association with metabolism and their hosts. We explored 471 metagenomes for viral populations from the pelagic epilimnion water column of temperate Lake Mendota. We unraveled the seasonal dynamics of viral populations and virus-host relationships at the genomic level. We examined abundance profiles, gene frequencies, and microdiversity patterns of viral populations over the 20-year time series to understand the evolutionary trajectories, adaptation mechanisms, interactions with microbial hosts, and influences of seasonality. We also studied the connections of viruses with environmental factors and posit that environmental constraints are mediated through virus-host interactions.

## Results

### Freshwater lakes harbor enormous uncharacterized viral diversity

In this study, we analyzed 471 metagenome samples covering six seasons each year^20^ (Fig. 1a, Table S1). These seasons—ice-on, spring, clearwater, early summer, late summer, and fall—are defined by environmental data and most accurately represent microbial phenology^20^. Our analysis identified a total of 1,820,639 viral scaffolds, with an average of approximately 2,600 viral scaffolds per metagenome (Fig. 1b, Table S2). Applying a stringent binning approach, we obtained 1,307,400 vMAGs (viral metagenome-assembled genomes or viral bins) (Fig. 1c, Fig. S1, and Table S3). In this study, a substantial number of viral genomes were generated, constituting approximately one-quarter of the entries within the IMG/VR v4 database, which encompasses around 5.6 million high-confidence viruses^3^.

**Fig. 1:**
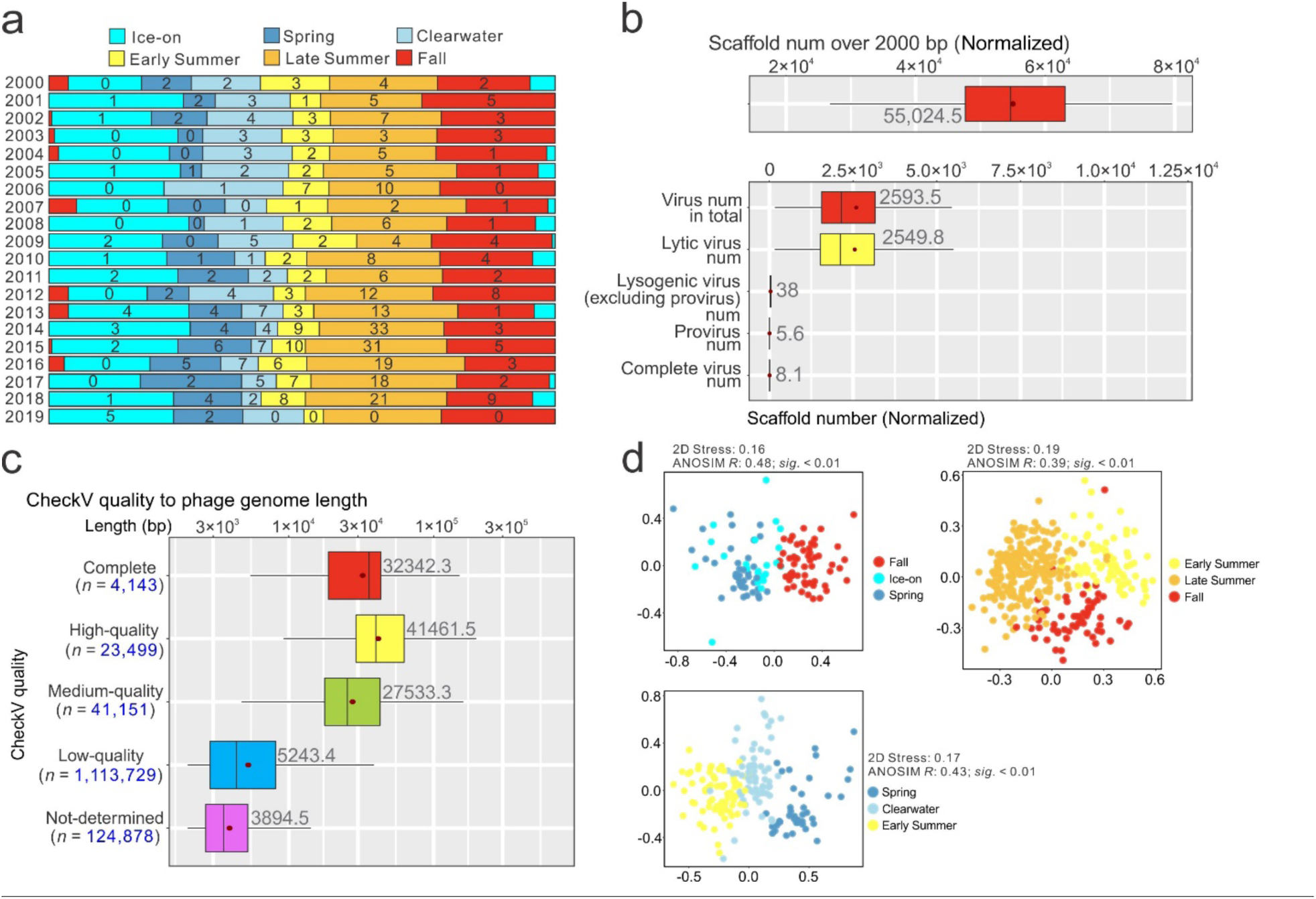
Statistics of viral scaffolds and genomes. **a** The number of metagenomes obtained in each season across 20 years. **b** Statistics of total metagenomic scaffolds and viral scaffolds across 471 metagenomes. Whiskers (outliers) are not shown for simplicity. The dark red points (with labeled values) indicate the mean values. Scaffold numbers were firstly normalized by 100M reads/metagenome to overcome uneven sequencing depth across samples. **c** Length and completeness of viruses after binning. CheckV quality to completeness range: Complete (100%); High-quality (90.0-100.0%); Medium-quality (50.0-89.99%); Low-quality (0.01-49.99%); Not-determined (labeled as “NA”). **d** NMDS plots presenting viral genome distribution among seasons. The viral genome abundances were calculated at the family level.

The use of viral genome binning significantly improved the length and completeness of our viral sequence collections, as assessed by CheckV^21^, leading to a notable enhancement in overall viral genome quality (Fig. 1c, Fig. S1, and Table S4). Furthermore, we clustered all vMAGs into 749,694 species based on 97% sequence identity. The number of species identified in our samples is comparable to approximately one-quarter of all virus species cataloged in the IMG/VR v4 database (high-confidence viruses) (749,694 vs 2,917,521), underscoring the substantial viral diversity in freshwater lakes. Notably, the rarefaction curve of species did not plateau, suggesting that there is still a considerable amount of unknown viral diversity in freshwater lakes (Fig. S1d). To delve deeper into the patterns observed, we examined the viral genome distribution at the family level, revealing distinct separation among viral communities across different seasons (Fig. 1d). Collectively, this not only highlights the richness of viral species but also emphasizes the seasonal and dynamic nature of viral communities.

Viral taxonomic classification revealed that dsDNA viruses in the class *Caudoviricetes* were the predominant group, followed by nucleocytoplasmic large DNA viruses in the class *Megaviricetes* (Fig. 2a). Most viral species identified within *Caudoviricetes* could not be taxonomically classified further, highlighting the need to investigate the extensive diversity of this class in freshwater environments (Fig. 2a). Less than 20% of the viral community received host prediction assignments (Fig. 2b). Across all six seasons, Bacteroidota were the most abundant host phylum, followed by Cyanobacteriota and Gammaproteobacteria (Fig. 2b). Each of the six assigned host phyla/classes displayed relative abundances of approximately 1-3% as revealed by the host MAG relative abundance distribution. The host prediction results revealed a diverse range of virus hosts, underscoring the extensive uncharted territory in this field (Fig. 2b).

**Fig. 2:**
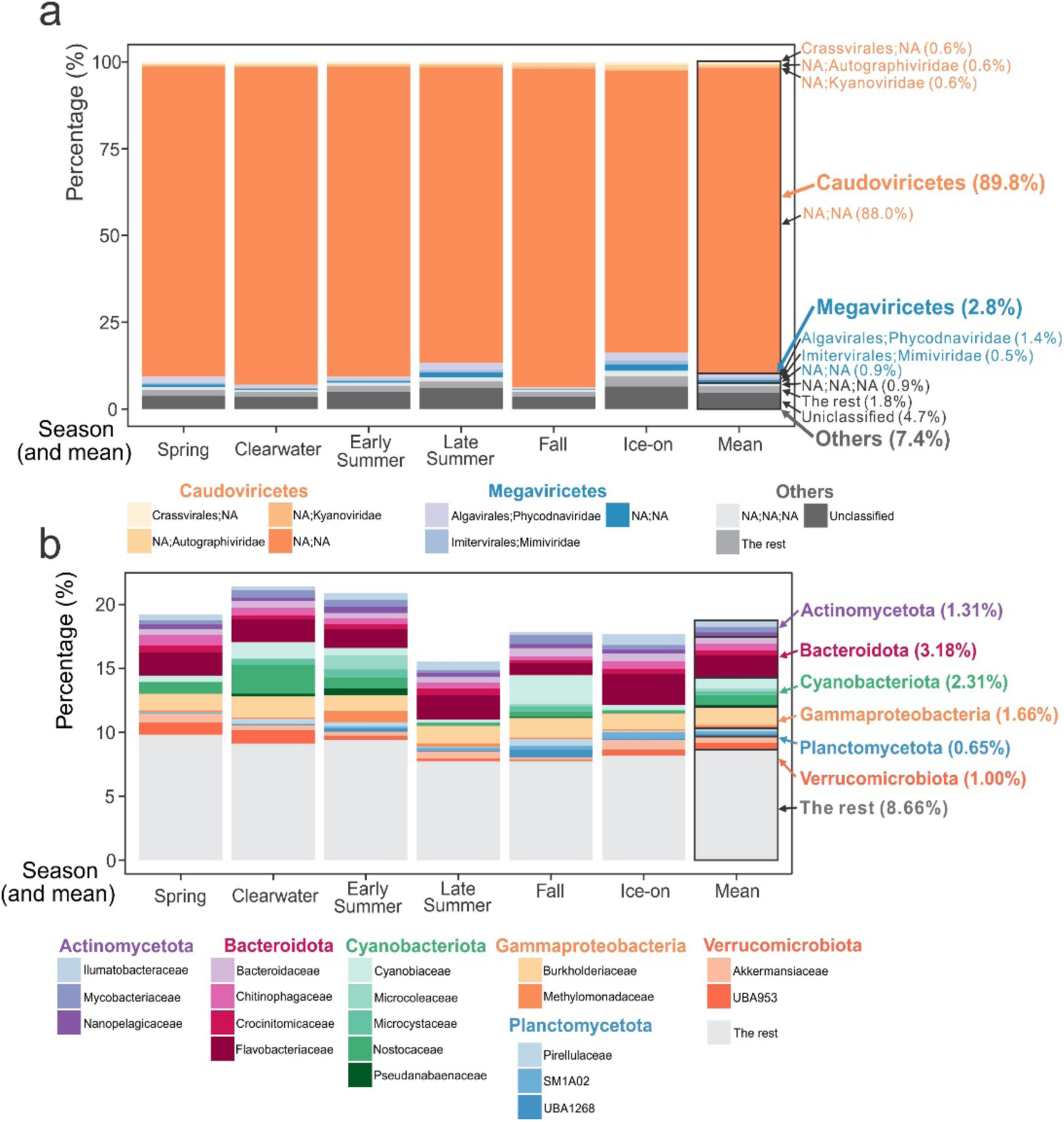
Seasonal abundance distribution of viruses and viral hosts. **a** Seasonal abundance distribution of viruses at the family level. Families of relative abundance < 0.5% across all seasons were combined as “The rest”. The “NA;NA” taxon within *Caudoviricetes* and *Megaviricetes* indicates families unclassified within the respective classes. Similarly, the “NA;NA;NA” taxon under “Others” represents other unclassified families. **b** Seasonal abundance distribution of viral hosts at the family level. Families of relative abundance < 0.5% across all seasons were combined as “The rest”. Unclassified host families were not depicted in the bar plot. Additionally, each bar plot includes a “Mean” bar, indicating the average percentage of relative abundances for all six seasons, with corresponding mean percentages clearly labeled in each plot.

### AMGs display distinct modes of distribution patterns: broad and narrow host ranges

We identified 150,458 AMGs from viral genomes, which clustered into 578 protein families (KEGG KOs, hereafter referred to as AMG clusters) (Table S5). These protein families demonstrated a diverse functional repertoire, comprising 12 distinct categories in total covering important biogeochemical transformations of C, N, and S (Fig. 3, Fig. S2, Table S6, and Supplementary Information). This repertoire was larger than those reported in previous studies (34 clusters discovered in the Global Ocean Survey communities and 322 clusters discovered in pelagic and benthic communities of the Baltic Sea)^22, 23^, highlighting the high diversity of our recovered vMAGs. The roles played by freshwater AMGs encompass crucial processes such as photosynthesis, methane oxidation, CO2 fixation for energy and carbon metabolisms, nitrogen metabolism for nucleotide biosynthesis, and sulfur metabolism for organosulfur degradation and sulfide production. We propose that, similar to other ecosystems, these AMGs likely provide substantial fitness benefits to viruses.

**Fig. 3:**
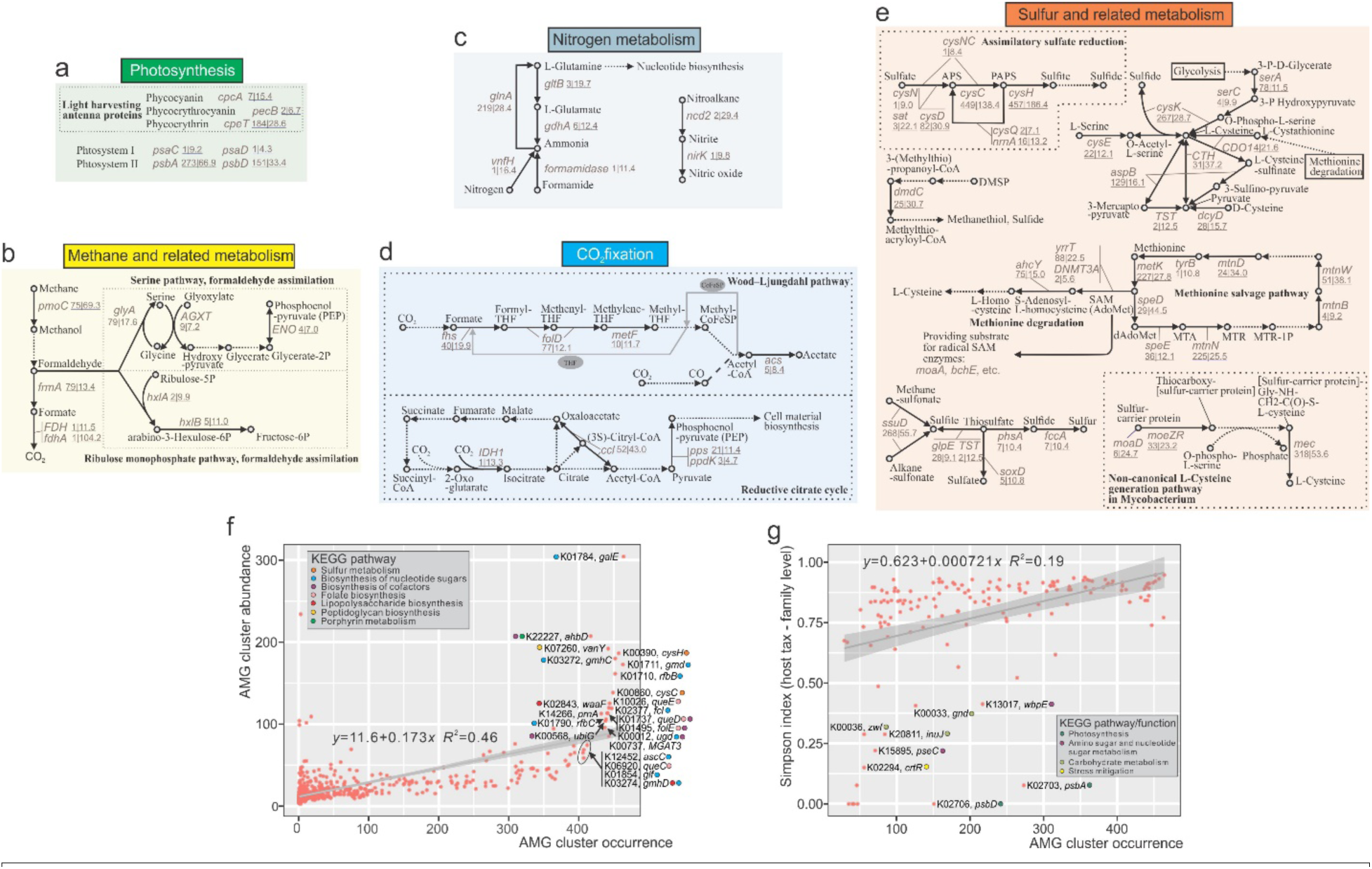
AMGs associated with carbon, nitrogen, and sulfur metabolism in freshwater viruses and modes of distribution patterns of AMGs. **a** AMGs involved in photosynthesis, **b** AMGs involved in methane and related metabolism, **c** AMGs involved in nitrogen metabolism, **d** AMGs involved in CO2 fixation, **e** AMGs involved in sulfur and related metabolism. Gene symbols are depicted in blue together with the occurrence and abundance values (labeled as “occurrence|abundance”; occurrence, the number of metagenomes in which an AMG cluster can be found; abundance, the mean normalized abundance of AMG cluster carrying viruses in the metagenomes that this AMG cluster can be found). Dotted arrows indicate steps that are not encoded by AMGs. Detailed information on each AMG cluster can be found in Table S6. Full AMG metabolisms and functions are depicted in Fig. S2. **f** The scatter plot of AMG cluster “abundance to occurrence”. High occurrence refers to distribution in > 400 metagenomes. AMG cluster KO dots are labeled with KO ID, gene name, and the corresponding KEGG pathway. AMG cluster abundance was normalized by 100M reads/metagenome. **g** The correlation of Simpson index of host taxonomy to AMG cluster occurrence. AMG clusters with low Simpson indices (narrow host range) as compared to the fitting curve are labeled with the KEGG pathway/function information (only AMG clusters distributed > 50 metagenomes were considered).

Two modes of viral auxiliary metabolism (namely broad and narrow) were discovered based on their distribution across broad or narrow host ranges, respectively. Broad host range AMGs were high-abundance AMG protein families that displayed a high occurrence distribution (identified in > 85% of all samples), were unaffected by seasonal change, and were resilient to intra-population dynamics of specific AMG-carrying viral species (Fig. 3f, Fig. S3). These AMG protein families were primarily involved in sulfur metabolism and cofactor and folate biosynthesis. The viral auxiliary metabolisms for organic and inorganic sulfur transformations often result in the production of sulfide as an end product^24^, which benefits host survival, growth, amino acid synthesis, protein function, and virion assembly. Additionally, the presence of cofactor and folate AMGs likely facilitate numerous reactions in energy production and stress alleviation (Supplementary Information).

Conversely, narrow host range AMGs revealed AMG protein families with limited host ranges that performed specific functions (Fig. 3g, Fig. S4). Examples of such specific AMG families include *psbA/D*, which encode photosystem II (PSII) reaction center domains D1/D2^8, 9, 11, 25^. These AMG families can enhance the photosynthetic activity of infected cyanobacteria. Previous research suggests that cyanophages can alter host carbon metabolism from carbon fixation towards the pentose phosphate pathway (PPP) for NADPH generation and deoxynucleotide biosynthesis^26^. In our analysis, we also identified specific AMG families associated with PPP, such as *gnd* and *zwf*^26^. Other examples include *crtG*, which is responsible for antioxidant production and mitigating stress from reactive oxygen species^27^ in *Burkholderiaceae*, thereby enhancing the overall fitness of virocells, and *inuJ* which encodes for the enzymatic degradation of sucrose into glucose in *Chitinophagaceae*, potentially serving as a means of energy preservation to support viral propagation. Overall, these two modes of AMGs highlight the adaptive strategies of viruses in their interactions with hosts and their significant involvement in microbially-mediated biogeochemical cycles.

### Seasonal patterns reveal tightly coupled virus-host dynamics and competition between viruses

Combining two decades of relative abundance data unveiled seasonal patterns of virus and host abundance. We examined three keystone microbial taxa: Cyanobacteria, Methanotrophs, and *Nanopelagicales* (ultrasmall acI within Actinobacteriota). In our analysis, we considered the abundances of both AMG-containing and non-AMG-containing viruses (Fig. 4). Among the 13 examined AMG clusters, when the viral genome completeness was high (75-100%), the majority of species members contained the corresponding AMG cluster KOs (> 85%) (Fig. 4a). This underscores the efficacy of delineating AMG-containing species representatives to represent all AMG-containing viruses.

**Fig. 4:**
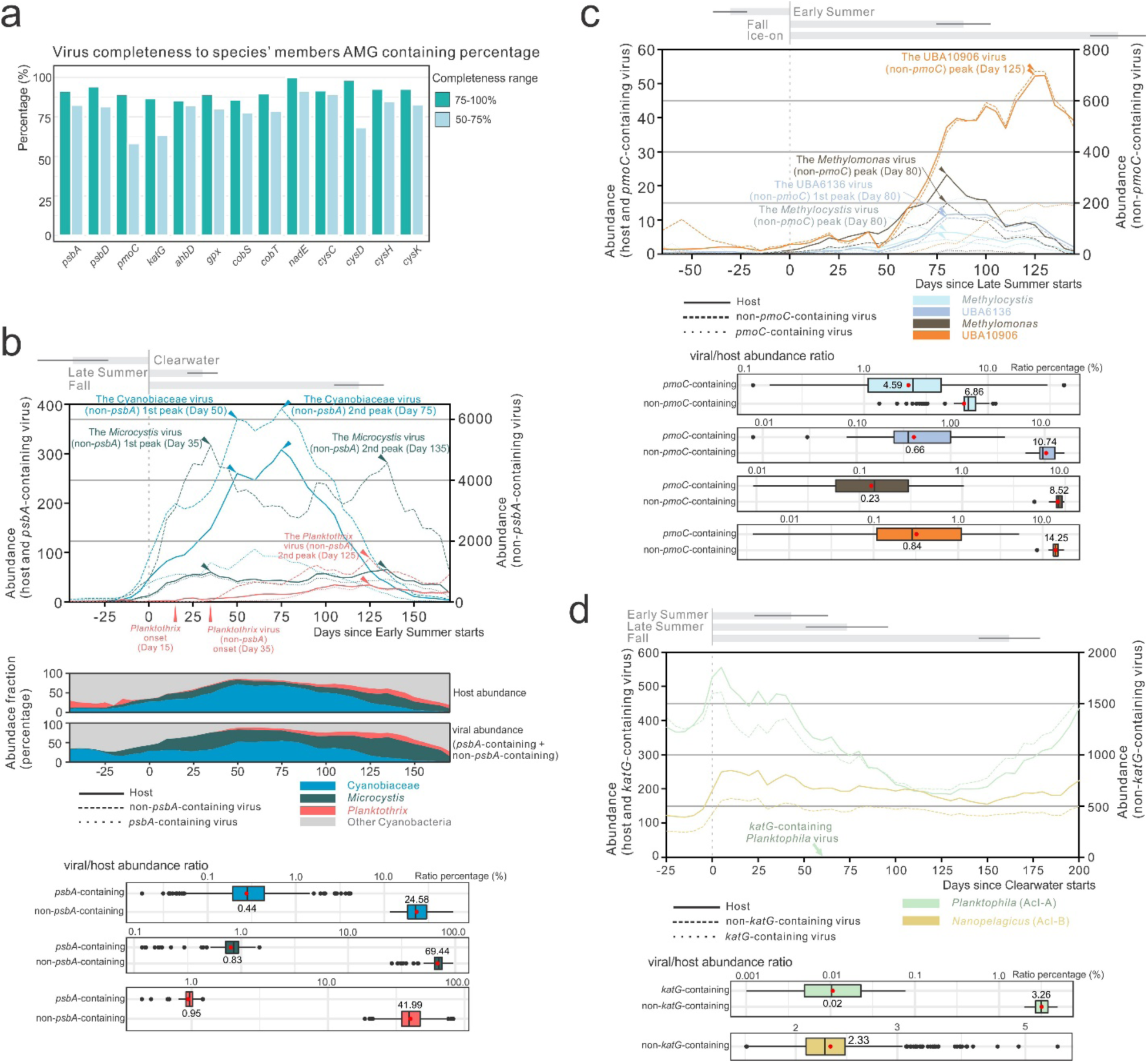
Viral-host correlation and seasonal change patterns for Cyanobacteria, methanotrophs, and Nanopelagicales. **a** Bar plot representing comparison of viral genome completeness to percentage of species’ members that contain AMGs. Species with representative genomes containing the corresponding AMGs were considered. **b** Seasonal change of viral and host abundances for Cyanobacteria groups. The first line chart depicts abundances of viruses and hosts of *Cyanobiaceae*, *Microcystis*, and *Planktothrix*. Individual lines represent mean abundances derived from interpolated values calculated at intervals of 5 days. The second chart depicts abundance fractions of *Cyanobiaceae*, *Microcystis*, *Planktothrix*, and other Cyanobacteria during seasonal change. The third box plot depicts viral/host abundance ratios of *Cyanobiaceae*, *Microcystis*, and *Planktothrix* across all timepoints. **c** Seasonal change of viral and host abundances for methanotroph genera. The first line chart depicts abundances of viruses and hosts of four methanotroph genera. The second chart depicts viral/host abundance ratios of four methanotroph genera. **d** Seasonal change of viral and host abundances for *Nanopelagicales* genera. The first line chart depicts abundances of viruses and hosts of two *Nanopelagicales* genera. The second chart depicts viral/host abundance ratios of two *Nanopelagicales* genera. The box plots on the top of each line chart of **b**, **c**, **d** indicate the means and standard deviations of the start day of corresponding seasons.

In three groups of Cyanobacteria, namely, *Planktothrix*, *Microcystis*, and *Cyanobiaceae*, host and virus abundances were positively correlated (Fig. 4b). Additionally, the peak abundance timepoints for these viruses (specifically, the non-AMG containing fractions) and hosts exhibited consistent patterns over two decades. In the case of *Planktothrix*, the virus onset date lagged the host onset date by ∼20 days, while the onset timepoints for viruses and hosts in *Microcystis* and *Cyanobiaceae* remained nearly identical (Fig. 4b). Previous research indicates host physiology and habitat controls influence viral progeny and suggests that fast-growing hosts could provide more resources for viral production^28, 29^. Therefore, one plausible explanation is that the *Planktothrix* virus population exhibits an extended lag phase, allowing for substantial replication when hosts achieve significant abundance levels at the start of the annual growth cycle. This may potentially constitute a viral propagation strategy specifically adapted for hosts with low abundance at the onset. *Microcystis* and *Planktothrix*, two genera potentially producing microcystin toxins^30^, reached peak abundances either at the beginning of late summer or fall, strategically avoiding the peak period of the most abundant *Cyanobiaceae* during late summer (Fig. 4b). This observation suggests a potential competition for the overlapping ecological niche, particularly concerning light and nutrient resources^31^, in which both viruses and hosts actively participate. This dynamic pattern provides valuable insights for studying the mechanisms underpinning niche competition and temporal succession, offering the potential to be employed as an approach for controlling harmful microcystin producing cyanobacterial groups in aquatic systems globally^32^.

Similar to patterns observed in Cyanobacteria, four methanotroph genera also had positive host-to-virus abundance correlations and seasonal abundance patterns (Fig. 4c). Notably, UBA10906 emerged as the most abundant genus, outcompeting the other three genera. Its peak abundance occurs later in the summer, extending into the fall and lasting longer compared to the other three genera. Specifically, we noted that the ratio of *Methylocystis* viruses containing *pmoC* (particulate methane monooxygenase subunit C) to their hosts was of a similar magnitude to the ratio of *Methylocystis* viruses lacking *pmoC* to hosts (Fig. 4c). In contrast, the comparisons involving the other three genera consistently showed that *pmoC*-containing viruses were less abundant by one order of magnitude than viruses lacking *pmoC*. Viral-encoded *pmoC* has the potential to augment aerobic methane oxidation^12^. An earlier study in soils showed that *Methylocystis* viruses were the most abundant ones receiving the CH4-derived carbon in soil microcosm incubations fueled by ^13^C-CH4^33^. We infer that an elevated abundance of viral *pmoC* can augment methane utilization in *Methylocystis* and fuel viral propagation.

Similarly, two *Nanopelagicales* (acI group) genera displayed a positive correlation between their host-to-virus abundance (Fig. 4d). The abundance of *Planktophila* during summer seasons is probably suppressed due to high concentrations of H2O2 produced by abiotic photochemical actions and biotic cyanobacterial and algal metabolisms which peak during this period^34^ (Fig. 4d). This suppression leads to a noticeable decline in *Planktophila* populations. While catalases encoded by *katG* are vital for reducing H2O2 levels and stabilizing *Planktophila* growth^35^, the low abundance of *katG*-containing viruses infecting *Planktophila* suggests they are insufficient in bolstering catalase activity to counteract the stress from elevated H2O2. Furthermore, heme is an essential cofactor in the generation of catalases. The *ahbD*-containing *Planktophila* virus abundance also represented a declining trend in late summer and fall (data not shown), indicating a constraint in virus-assisted heme synthesis for catalase production. Consequently, despite the presence of these viruses, the high H2O2 concentrations during summer likely contribute significantly to the observed decline in *Planktophila* abundance.

Among these three host groups with significant biogeochemical roles, the ratios of AMG-containing viruses to hosts were consistently one to two orders of magnitude lower than those of non-AMG-containing viruses to hosts. Considering that most previous studies did not enumerate the abundance of AMG-containing viruses in nature, there might have been an excessive emphasis on the significance of viral AMGs associated with the metabolism of specific substrates (such as *pmoC* for methanotrophs in methane utilization^12^) or enhancing rate-limiting enzymes (such as *psbA* for Cyanobacteria to optimize photosynthesis^9, 11^). Furthermore, we propose that the belief that AMG-containing viruses are more prevalent and important in the community primarily stems from isolated strains. For instance, almost all Myoviruses and over half of Podoviruses infecting Cyanobacteria are believed to have *psbA* in their genomes^36^. However, based on the findings of this study, observations of relative abundances suggest that non-AMG-containing viruses overwhelmingly prevailed and closely mirrored the seasonal fluctuations of their hosts. This implies that non-AMG-containing viruses make up the majority of viral communities. Overall, we propose that the magnitude of influence of AMGs on viral fitness, metabolism, ecosystem function, biogeochemistry, and their adaptations to hijacking hosts likely needs reevaluation in future research^37^. Furthermore, comprehensive studies are essential to explore the interactions between non-AMG-containing viruses and hosts thoroughly.

### Persistent viral populations exhibit intra-population diversity undergoing evolutionary progressions and soft genome-wide sweeps

To examine intra-population diversity among persistent viral species, we studied viral species with AMGs encoding four important functions (*psbA* for photosynthesis, *pmoC* for methane oxidation, *katG* for reducing H2O2 stress, and *ahbD* for heme synthesis) (Table S7 and Supplementary Information). In our analysis of the 471 metagenomes, four of the six *psbA*-containing viral species demonstrated a positive correlation between species abundance and nucleotide diversity (*p*-value < 0.05) (Table S8). Moreover, two *ahbD*-containing viral species characterized by high occurrence showed a positive correlation between species abundance and SNP density (*p*-value < 0.05) (Table S8).

We subsequently expanded our analysis to encompass all viral species containing AMGs. Of these, 221 out of 866 species with valid nucleotide diversity results and 262 out of 777 species with valid SNP density results showed significant positive correlations with viral abundance, respectively. In contrast, only 23 out of 866 species with valid nucleotide diversity results and 12 out of 777 species with valid SNP density results showed significant negative correlations with viral abundance, respectively (Table S8). These findings suggest that viral intra-population diversity is mainly governed by the neutral theory^38^, wherein an augmented population size generally leads to increased nucleotide diversity and SNP density.

Positively selected genes within these persistent viral populations (Table S9) encoded enzymes associated with purine biosynthesis, viral RNA synthesis, DNA repair, controlling of cellular and viral DNA and mRNA turnover, transcriptional regulation, bacterial cell wall penetration, as well as auxiliary metabolisms related to photosynthesis (*psbA*), heme synthesis (*ahbD*), and folate biosynthesis (*moaA*). Such findings suggest a mechanism of viral fitness selection associated with viral infection and host regulation, virion replication, and host metabolism redirection or augmentation^1, 10, 39^.

In certain viral species, whole genome genetic heterogeneity gradually decreased (Fig. 5a), as evidenced by the fact that persistent *psbA*-and *ahbD*-containing viral species exhibited an increasing SNP allele frequency over time, as demonstrated by linear regression (high regression slope) and Spearman’s rank correlation tests (significant *p*-value) (Fig. 5a, Table S11). We examined the genes containing these increasing-frequency SNP alleles (Fig. 5a). Six of the nine viral genes were positively selected (Table S9) within these two vMAGs; three were annotated with important functions (*psbA* and *ahbD* for auxiliary metabolisms, restriction endonuclease type II-like genes for host genome degradation, nucleotide recycling for viral replication^40^, and exclusion of superinfections^41^), contributing to viral fitness. This indicates that, while these high occurrence viral species persist over time, their sub-populations have changed. Specifically, selection favored some sub-populations with advantageous alleles. Nevertheless, the mean allele frequencies across genomes remain relatively low (∼0.7) (Fig. 5a), implying that either the genome-wide sweep remains ongoing or viral populations undergo a “soft sweep”, wherein selection favored a few sub-populations from large, diverse populations^42, 43, 44^. Bacteriophages typically have higher genomic diversity and recombination rates than bacteria^45, 46^. Due to the high microdiversity that existed prior to the start of this study, it will take a longer time for sub-populations with selection advantages to take over the population. Concurrently, an elevated recombination rate seems to counteract selection, promoting recombination amongst sub-populations with distinct micro-niches, thereby preserving genome-wide diversity.

**Fig. 5:**
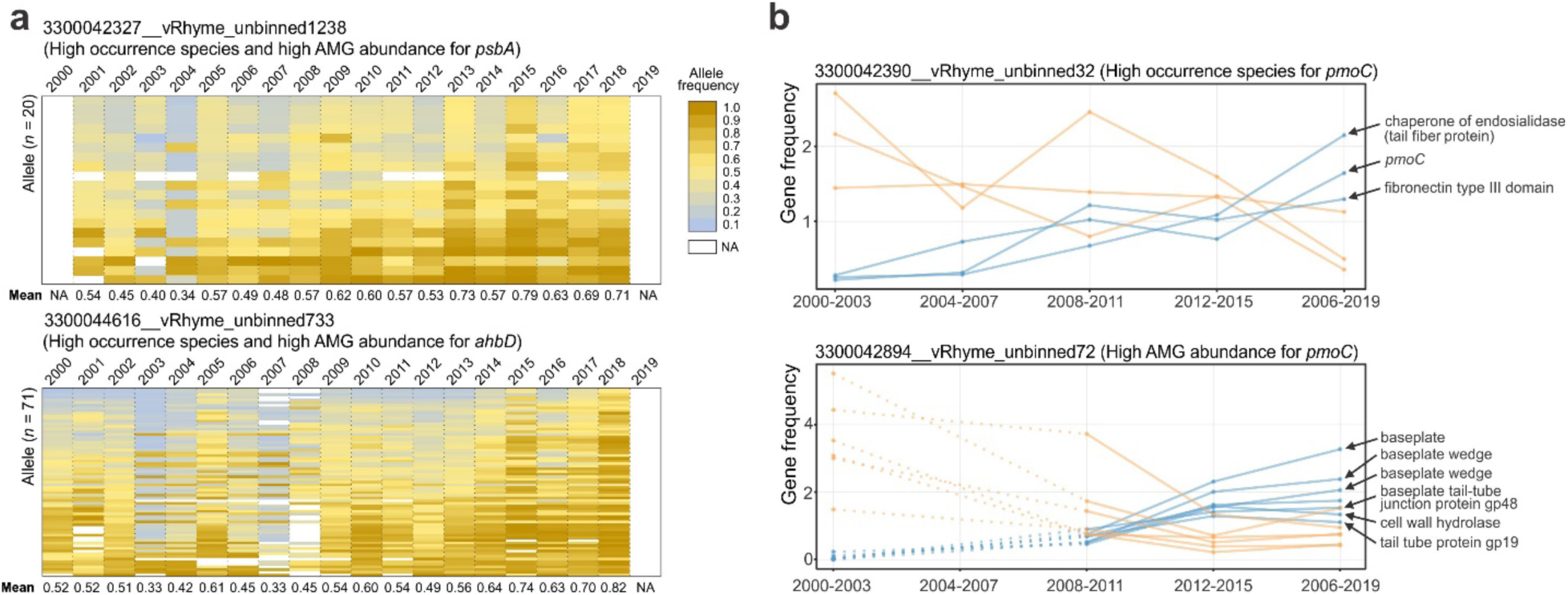
Microdiversity changes of persistent viral populations. **a** Pattern of SNP allele frequency variation. SNP alleles are organized in ascending order based on the mean allele frequencies over 20 years, with each row denoting an individual SNP. The mean allele frequencies across the genome are labeled below each column accordingly. The allele frequency is depicted by the proportion of reads aligning to the reference allele – the predominant allele in the corresponding viral genome for 2018. **b** Gene frequency change pattern. The y-axis represents gene frequency, determined by dividing gene coverage by the average coverage of all other genes within the genome. Genes exhibiting a mean frequency change of ≥ 1.0 between 2000-2003 and 2016-2019 are considered to have significantly increased or decreased in frequency. Intervals devoid of meaningful gene frequency values are represented with dash lines in the graph.

Additionally, for some viral populations, certain genes have either increased or decreased gene frequencies over time (Fig. 5b, Table S12). The increasing-frequency gene repertoire encodes structural proteins, such as chaperones of endosialidase (tail fiber proteins for initial absorption of virus into the host^47^), baseplate, baseplate wedge, and tail tube proteins; viral core function proteins, such as fibronectin type III containing protein (probably for virus-cell surface interaction^48^) and cell wall hydrolase (for host cell wall degradation and facilitating bacteriolysis and virion release^49^); and an AMG protein (PmoC). This indicates the importance of virus structural proteins, viral infection proteins, and auxiliary metabolic proteins in strengthening viral fitness. This scenario suggests a similar genome-wide selection pattern in that certain sub-populations that harbored important functional genes in the lake prior to this study (before 2000) gradually became dominant in the populations from 2000 to 2019. Collectively, despite viral populations having a high level of diversity and rate of recombination, selections for genes with fitness advantages and genome-wide selections still play an important role in viral population dynamics.

### Environmental constraints indirectly influence viral abundance via “top-down” and “bottom-up” controls

Host dynamics are controlled by both top-down (e.g. grazing by protists, viral lysis) and bottom-up (e.g. water temperature, nutrient concentrations) drivers^4^. We expect these dynamics to also manifest in measured viral abundances and potential viral roles. We focused on Cyanobacteria and their viruses to explore whether available limnological measurements could explain their dynamics. The number of duration days in which the Cyanobacteria and Cyanobacteria virus abundances were > 20% of their peak abundances were related to the environmental parameters using Spearman’s rank correlation test. The number of duration days should reflect an integrated influence of the environmental conditions during the summer season. As expected, water temperature and Secchi depth (a measure of water clarity) were positively correlated with both Cyanobacteria and their viruses (Fig. 6a and Table S13).

**Fig. 6:**
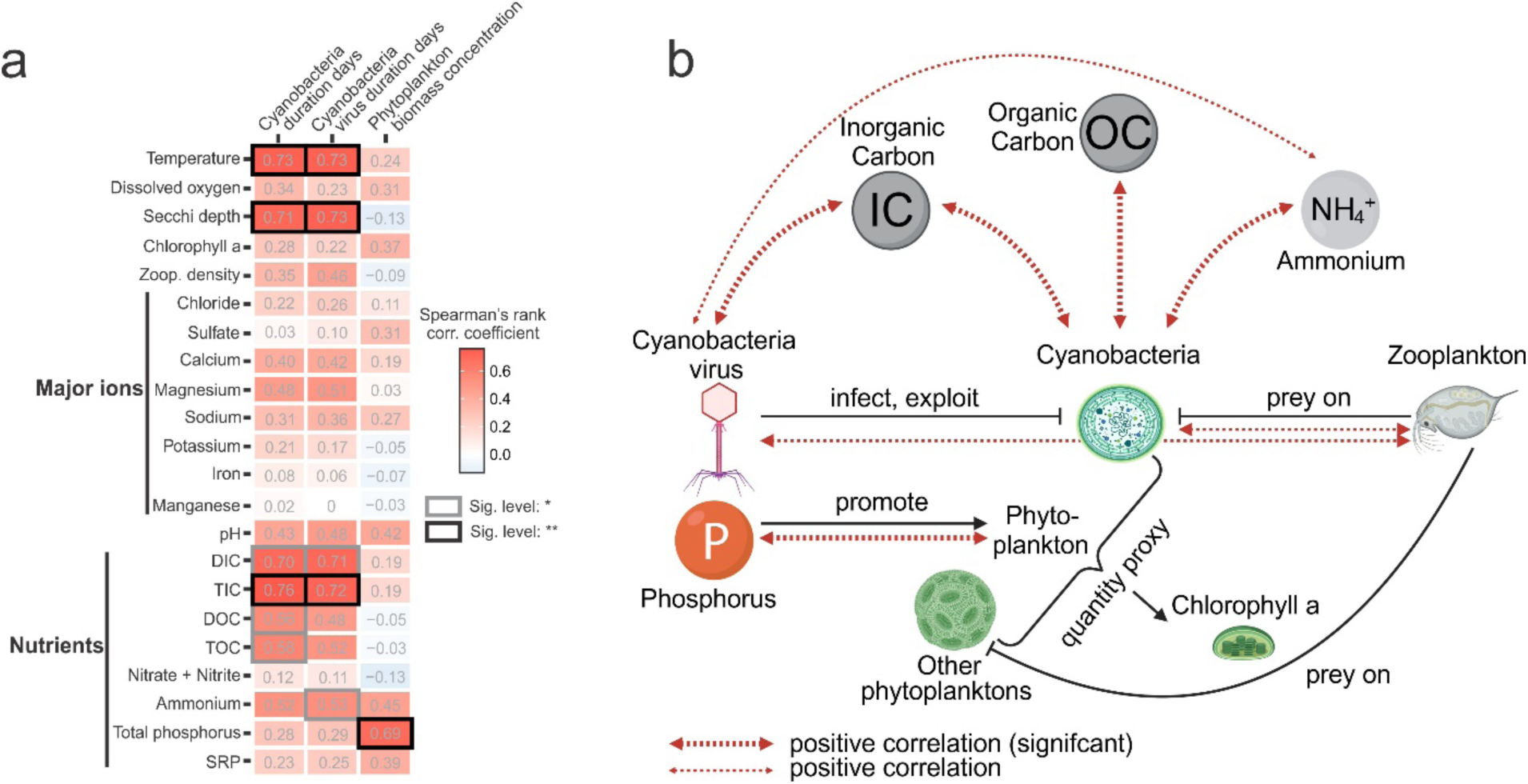
Correlations between viruses, hosts, and environmental parameters across the time-series. **a** Heatmap representing the Spearman’s rank correlation between environmental parameters and cyanobacteria, cyanobacteria virus, phytoplankton biomass concentration, and zooplankton density. Spearman’s rank correlation coefficients are labeled in individual cells and significance levels are indicated by grid borders, with black borders denoting *p*-values < 0.01 (labeled as “**”) and grey borders denoting *p*-values < 0.05 (labeled as “*”). **b** The schematic diagram shows the interactive connections among the environment, viruses, bacteria, phytoplankton, and zooplankton.

These relationships reflect the intricate balance of both “top-down” and “bottom-up” factors that shape the environment, viruses, bacteria, and predators within the ecosystem (Fig. 6 and Table S13). Phosphorus, acting as a bottom-up factor, stimulates phytoplankton growth^50^. This increase in phytoplankton biomass forms the basis for further ecological interactions. Inorganic carbon serves as the primary carbon source for cyanobacteria through photoautotrophy, promoting both cyanobacteria and viral proliferation in a bottom-up manner. Additionally, several cyanobacteria can assimilate organic carbon concurrently during photosynthesis, and the mixotrophic metabolism accelerates the growth of Cyanobacteria^51, 52^. This aligns with the observed positive correlation between organic carbon and Cyanobacteria (Fig. 6b). Ammonium also correlated positively with cyanobacterial virus abundance (Fig. 6a). As the primary nitrogen source for assimilation by Cyanobacteria^54^, it promotes Cyanobacteria growth, which in turn, supports the proliferation of Cyanobacteria viruses (Fig. 6b). Conversely, reflecting a “top-down” control, as Cyanobacteria abundance increases, zooplankton density increases, demonstrating a typical predator-prey dynamic. The increased cyanobacterial abundance provides more hosts for cyanobacteria viruses, which leads to the elevation of virus abundance as evident from the observed correlations in virus-to-host abundance (Fig. 6b). These “top-down” interactions are critical in regulating the populations within the ecosystem.

## Discussion

Time-series studies at the fine-scale resolution of viral genomics can resolve the role of viruses in nutrient, biogeochemical, and energy transformations via virus-host interactions and provide a more holistic context in microbiomes and ecosystems. In this study, we used metagenomics on samples from a 20-year time-series to unravel and characterize viral diversity, ecology, and evolution, and the metabolic processes viruses control and impact in a freshwater lake. We uncovered unprecedented viral diversity from freshwater systems over a 20-year time-series. Our study highlights the enormous volume of unknown viral diversity found in a single temperate freshwater lake, which suggests even more around the world, and additionally suggests that other freshwater systems, such as tropical lakes, are likely purveyors of viruses playing important roles in nutrient and biogeochemical transformations that require further investigation.

Our study also provides viral AMG collections that are significantly larger than previous reports^22^ in terms of both gene families and their functions. In this study, two modes of AMGs (broad vs narrow) were described. On the one hand, broad AMG clusters tend to have high abundance in individual lake samples, high occurrence across the time-series, widely prevalent distribution despite seasonal changes, and resilience to intra-population dynamics of specific AMG-carrying viral species. These AMG clusters are primarily involved in sulfur metabolism and cofactor and folate biosynthesis. Viral organo/inorganic-sulfur auxiliary metabolism can help assimilate sulfate and produce sulfide as end products^24^, widely benefiting host survival/growth, amino acid synthesis and protein function, and virion assembly. Cofactors and folates can facilitate many reactions in energy production and stress alleviation. These broad host range AMGs represent a repertoire of universal functions interacting with a range of hosts for enhancing viral fitness. On the other hand, as reflected by specific AMG cluster-to-host connections, narrow host range AMGs have specific functions for enhancing viral fitness. These narrow range AMGs exhibit targeted adaptive mechanisms that enhance viral fitness through interactions with their hosts and undergo seasonal dynamics in parallel with those of their hosts. The two AMG modes highlight the adaptive strategies of viruses in interacting with hosts and their significant involvement in microbially-mediated biogeochemical cycles. Collectively, viral auxiliary metabolisms as reflected in this study, reveal an unprecedented and comprehensive functional repertoire encompassing 12 distinct categories (for details, see Supplementary Information), emphasizing the need for additional research to uncover viral-host interactions and the underlying ecological significance of AMGs.

Few studies have focused on the evolution and environmental analyses of viral population dynamics, particularly for a two-decade time-series study of the natural environment. In this study, persistently distributed viral populations of high occurrence underwent both positive gene selection and genome-wide selection. Three evolutionary processes were observed: selection favored genes associated with fitness, genomic heterogeneity decreased over time, and sub-populations carrying certain genes became dominant. Similar to a lake green sulfur bacterial population in which SNP variations were slowly purged and some genes were either swept through or lost within the population over time^6^, our study indicates the universality of evolutionary processes in both viruses and microorganisms. The selected viral genes mostly augment viral fitness by facilitating genome replication, regulating host transcription, releasing stress effects on the host, and providing auxiliary functions to improve the overall fitness of virocells^24^. These processes can be jointly explained by the concept that some sub-populations with advantageous traits acquired through mutations or horizontal gene transfer, outcompete others and become predominant in the observed populations^6, 42, 43^, which appear to be “stable” when only viewed from a macrodiversity perspective.

In the evolutionary arms race between viruses and their hosts, “kill-the-winner” and other forms of dynamics frequently occur, causing fluctuations in the abundance of various viral strains^55^. Despite these fluctuations, certain viral species persist over extended periods and demonstrate high occurrence over time, indicating their evolutionary success in adapting to changing environmental conditions. These high occurrence viral species may represent a ‘royal family’ viral species in the model used to explain the “kill-the-winner” dynamics^55^, where certain sub-populations with enhanced viral fitness have descendants that become dominant in subsequent “kill-the-winner” cycles. It is probable that these high occurrence viral species maintain a stable presence at the coarse diversity level while undergoing continuous genomic and physiological changes at the microdiversity level. The dynamics at the level of viral and host interactions play a pivotal role in driving viral evolution and maintaining the dominance of ‘royal family’ viral species. For example, the selection of viral genes associated with resistance and counter-resistance results in enhanced bacterial cell wall penetration, initial absorption into the host cell, and virus-cell surface interactions. The selection of viral genes associated with nucleotide biosynthesis and recycling facilitates more efficient viral replication by better exploiting host resources. Finally, the selection and increased frequency of AMGs such as in in carbon, nitrogen, and sulfur metabolisms alleviate host metabolic bottlenecks, and better exploit metabolic substrates and energy from the environment. Therefore, the sustained interactions and co-evolution of viruses and hosts over time drive suggest better adaptation of highly abundant viral species to local environmental conditions.

Concurrently, we discovered that environmental factors, such as inorganic carbon and ammonium indirectly influence viral abundance through virus-host interactions. To our knowledge, this study represents the first statistical analysis linking viral abundance to environmental factors through time-series datasets. Our observations suggest a complex interplay of “bottom-up” controls, such as nutrient availability and primary production, and “top-down” controls like predator-prey dynamics. These analyses provide valuable insights into the vast viral diversity, ecological and evolutionary dynamics, and the roles of viruses in biogeochemistry in globally distributed freshwater systems. Overall, our findings underscore the necessity for further research on viruses in microbiomes and ecosystems, and for a holistic approach that places viral studies in the broader context of biodiversity, virus-host interactions, and the physico-chemical constraints existing in natural environments.

## Reporting Summary

Further information on research design is available in the Nature Research Reporting Summary linked to this article.

## Data availability

The metagenomic datasets (including assemblies and raw reads) are all available under JGI Proposal ID 504350 at the platform of the Integrated Microbial Genomes & Microbiomes system (IMG/M: https://img.jgi.doe.gov/m/). The retrieved viral genomes were deposited in NCBI Bioproject X. At the same time, the TYMEFLIES viral genomes and related properties, including annotations for viral proteins, taxonomic classification, host prediction, and virus clustering results, were deposited in the following address: https://figshare.com/articles/dataset/TYMEFLIES_vMAGs_and_related_properties/24915750.

The raw environmental parameter spreadsheets are available in the Environmental Data Initiative (EDI, https://edirepository.org/) database.

## Code availability

Codes used in this project are available at the following GitHub repository: https://github.com/AnantharamanLab/TYMEFLIES_Viral

## Acknowledgments

We thank the local support for fieldwork conducted in Lake Mendota, WI as a site of a long-term lake ecological study for North Temperate Lakes Long Term Ecological Research, including the following people as sampling leads: Angela Kent, Tony Yannarell, Ashley Shade, Stuart Jones, Ryan Newton, Georgia Wolfe, Emily Kara Read, Lucas Beversdorf, James Mutschler, and Robin Rohwer; and initial Microbial Observatory lead Eric W. Triplett. This research was supported by the National Science Foundation grant number DBI2047598 (KA), USDA National Institute of Food and Agriculture under Hatch project 1025641 (KA), and Simons Foundation Investigator in Aquatic Microbial Ecology Award LI-SIAME-00002001 (BJB). CM was supported by a National Science Foundation Graduate Research Fellowship. RRR was supported by a National Science Foundation Postdoctoral Research Fellowship in Biology (DBI-2011002). Sequencing and initial sequencing datasets processing were carried out at the U.S. Department of Energy Joint Genome Institute (CSP 504350). The work (proposal: CSP 504350) conducted by the U.S. Department of Energy Joint Genome Institute (https://ror.org/04xm1d337), a DOE Office of Science User Facility, is supported by the Office of Science of the U.S. Department of Energy operated under Contract No. DE-AC02-05CH11231.

## Author contributions

ZZ, KDM, and KA conceived the project. RRR performed DNA extraction and sequencing. ZZ, CM, and PQT conducted bioinformatic analyses, statistical analyses, visualization of results, and content organization. ZZ and KA wrote the manuscript draft. All authors (ZZ, KDM, KA, RRR, BJB, CM, PQT) reviewed the results, edited, and approved the manuscript.

## Methods and Materials

### Samples

In this study, 471 water filter samples were collected from a pelagic integrated 12-meter zone in Lake Mendota, Madison, WI, USA (GPS: 43.0995, -89.4045). Lake Mendota is a eutrophic freshwater lake located in Madison, WI (size: 39.4 km^2^; average depth: 12.8 m; pH: 8.5) and an important component within the North Temperate Lakes Long Term Ecological Research project (NTL-LTER) started in 1981^6^. The samples were collected over several timepoints across different seasons each year, and the total sample period spanned 20 years (2000-2019). For each sample date, an approximately 150 mL integrated water sample was collected by filtering through a 0.2 μm pore size polyethersulfone Supor filter (Pall Corporation, Port Washington, NY)^56^. Filters were stored at −80°C for long-term storage. For omics sequencing, DNA extraction was conducted by using the FastDNA Spin Kit (MP Biomedicals, Burlingame, CA) with minor modifications^57^.

### Environmental parameters

Environmental parameters, such as temperature, dissolved oxygen, Secchi depth, chlorophyll a, major ions, limnological nutrients, phytoplankton biomass concentration, and zooplankton density, were acquired from the sampling station at Lake Mendota (GPS: 43.0988, −89.4054) through the NTL-LTER program (https://lter.limnology.wisc.edu/) or measurements conducted by the McMahon Lab. The original datasets were processed and organized from the Environmental Data Initiative (EDI, https://edirepository.org/) database with the following identifiers: chlorophyll a (knb-lter-ntl.38.28), major ions (knb-lter-ntl.2.38), limnological nutrients (knb-lter-ntl.1.60), phytoplankton biomass concentration (knb-lter-ntl.88.31), and zooplankton density (knb-lter-ntl.90.33). Temperature, dissolved oxygen, and Secchi depth datasets were collected, processed, and integrated from diverse sources, including EDI data of knb-lter-ntl.335.1, knb-lter-ntl.29.29, knb-lter-ntl.130.29, knb-lter-ntl.400.2, knb-lter-ntl.129.31, knb-lter-ntl.31.30, and McMahon Lab manual profiles (Year 2006-2012 McMahon Lab and Year 2013-2019 Rohwer Team profiles).

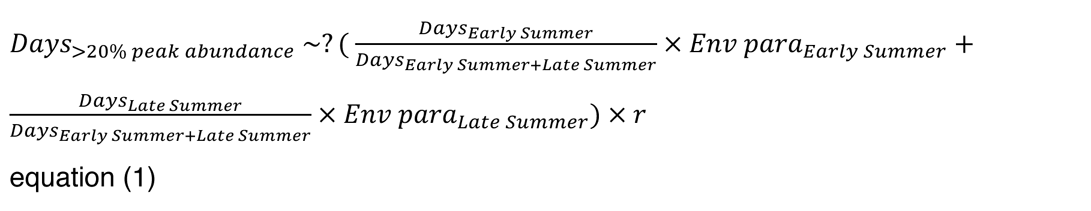

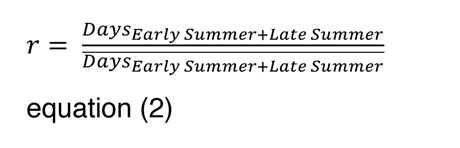

The two equations presented above show the method to assess potential correlations between the duration days in which the abundance is maintained at > 20% of the peak abundance (for both cyanobacteria and cyanobacteria viruses) and the average environmental parameters throughout summer seasons. These environmental parameters were obtained from both Early Summer and Late Summer, and their significance was weighted based on the dates of each summer season. In equation (2), “r” represents the ratio employed to standardize the summer dates for each year in relation to the mean summer dates. Duration days were determined by computing the abundance profile using an interpolation function for each year, with intervals of five days. The interpolation function was applied to abundance data within the time range of -45 to 160 days (since the start of Early Summer for each year). The years that could not meet the entire time range were excluded from the correlation analysis. The Spearman’s rank correlation test with Fisher z-transformation was employed, and the *p*-value was provided. The mean value of each parameter for every season was calculated by averaging all the measurements within that specific season (some missing values were denoted as “NA”).

### Metagenome sequencing and processing

Extracted DNA from 471 samples was submitted to the Department of Energy Joint Genome Institute (DOE JGI) (Walnut Creek, CA) for metagenomic sequencing. Illumina regular fragments with ∼300 bp length were made for metagenome library construction; afterward, the high-throughput sequencing was conducted using the Illumina NovaSeq S4 (Illumina, San Diego, CA), yielding paired-end reads of 150 bp each and approximately 1.5×10^8^ reads (accounting for both ends) per sample. The metagenome assembly was performed by metaSPAdes v.3.14.1^58^ and annotated by the IMG annotation pipeline (IMGAP) v5.0.20^59^ implemented in the Integrated Microbial Genomes & Microbiomes system (IMG/M: https://img.jgi.doe.gov/m/).

### Virus identification and genome binning

VIBRANT v1.2.1^60^ was used to identify and annotate virus (bacteriophage) scaffolds from metagenomic assemblies with default settings. Only viral scaffolds (including both provirus and non-provirus) longer than 2000 bp were used for downstream analysis. vRhyme v1.0.0^61^ was used to reconstruct viral metagenome-assembled genomes (vMAGs) from the identified viral scaffolds with default settings. The following four criteria were used to refine the best vMAG (or bin) collection suggested by the default result of vRhyme: 1) Proviruses identified by VIBRANT were excluded from binning. 2) Two or more lysogenic (non-provirus) viral scaffolds cannot be in the same bin. 3) Viral scaffolds identified by CheckV v0.8.1^21^ (database checkv-db-v0.6) as “Complete” were excluded from binning. 4) The maximum number of bin redundancy should be ≤ 1. Any predicted bins that did not meet the above four criteria were split into individual viral scaffolds. CheckV was also used to estimate the vMAG quality with default settings. Since CheckV can only process single-contig viral genomes, for each vMAG, we first linked vMAG scaffolds with 1500 ‘N’s to make temporary ‘single-contig’ viral genomes.

### Virus clustering

We first clustered all viral genomes into families and genera using the gene sharing and AAI method, as described in Nayfach et al.’s previous publication^62^. An all-vs-all DIAMOND BLASTP (v0.9.14.115) was performed for all virus genome protein sequences with the settings of “--evalue 1e-5 --max-target-seqs 10000 --query-cover 50 --subject-cover 50”^62^. Then, the gene sharing numbers between each pair of genomes, as well as the average amino acid identity of shared proteins, were parsed from DIAMOND BLASTP results. Edges (based on the minimum values of gene sharing and AAI) and nodes (viral genomes) were parsed accordingly for both families and genera, then filtered and subjected to MCL-based^63^ (v14.137) network clustering. The edge filtering criteria and settings of the MCL inflation factor were adopted from the previous publication^62^ (https://github.com/snayfach/MGV/tree/master/aai_cluster).

For non-singleton genera, we further clustered them into species. dRep v3.2.2^64^ was used for dereplicating all viruses within each genus with the settings of “-l 2000 --ignoreGenomeQuality - pa 0.8 -sa 0.95 -nc 0.85 -comW 0 -conW 0 -strW 0 -N50W 0 -sizeW 1 -centW 0”. The resulting representatives (the best representative viral genomes picked according to genome length) together with singleton genera and individual viral genomes that were not assigned to any genera were the final collection of species.

### Taxonomic classification

We combined three approaches to conduct taxonomic classification. For the first and second approaches, we adopted the procedure as described in the instructions as suggested previously^3^. For searching against NCBI RefSeq viral proteins, DIAMOND BLASTP v0.9.14.115 was used to BLAST against NCBI RefSeq viral proteins (2023-01-13 release)^65^ using all the TYMEFLIES viral proteins with settings of “blastp -evalue 1e-5 --query-cover 50 --subject-cover 50 -k 10000”. For any viral genome with ≥ 30% of proteins having significant hits to NCBI RefSeq viral proteins, a ≥ 50% majority taxonomy was assigned based on the taxonomy (using reformatted ICTV taxonomy with eight ranks) of the best hits of individual proteins. For the VOG marker HMM searching approach, hmmsearch (HMMER v3.1b2^66^) was used to search against VOG database v97 (2021-04-19 release, http://vogdb.org) using all the TYMEFLIES viral proteins. Only 587 VOG marker HMM profiles were used as the reference for taxonomic classification^3^. The criteria for positive hits were score ≥ 40 and E-value < 1e-5. The taxonomy of a viral genome was obtained based on individual markers detected using a simple plurality rule if multiple hits were present. For the third approach, geNomad v1.5.1 was used to annotate viruses and get taxonomy from the annotation result using default settings^67^. For each vMAG, we first linked vMAG scaffolds with 1000 ‘N’s to make temporary ‘single-contig’ viral genomes in order to meet the input requirement of geNomad.

If a viral genome was not assigned taxonomy by any of the above three approaches while it was placed in a genus with the other member(s) assigned using the NCBI RefSeq viral protein searching approach, the lowest common ancestor (LCA) of this genus was used as the taxonomic classification (in this case, the deepest LCA rank is limited to the genus level). All the aforementioned taxonomic classification approaches were labeled in accordance with the taxonomy obtained. If overlaps occurred, the top-order approach was given the highest priority.

### Host prediction

We used three approaches for predicting the hosts of viruses. For the first approach, iPHoP v1.2.0^68^ was used to predict the host from all viruses (‘N’-linked sequences to make temporary ‘single-contig’ viral genomes) using the default settings. The TYMEFLIES species representative metagenome-assembled genomes (2855 MAGs total, dereplicated by dRep^64^ with 96% sequence identity cutoff) were added to the default iPHoP database “Sept_21_pub”. The host prediction to genome results (based on host-based tools, including “blast”, “CRISPR”, and “iPHoP-RF” results) were finally assigned with the following rules: 1) If “blast” or “CRISPR” results were obtained for one virus genome, the result with the highest confidence score was assigned as the final result. 2) If only “iPHoP-RF” results were obtained for one virus genome, the result with the highest confidence score was assigned as the final result. For the second approach, we predicted the viral host based on the auxiliary metabolic gene (AMG) identity match between AMGs and microbial counterpart genes. The viral AMGs were identified, filtered, and summarized (details refer to the following section). The counterpart genes (with the same KEGG Orthology) were parsed out from all TYMEFLIES MAGs. For any viruses that have AMGs connected to their counterparts in TYMEFLIES MAGs (potential viral contigs were excluded as mentioned above) based on the cutoff of sequence identity ≥ 60% (with DIAMOND BLASTP options of “--query-cover 70 --subject-cover 70”), the AMG viral-host connections were established. Similarly, such multiple viral-host connections based on AMGs for a virus were aggregated and the lowest rank with ≥ 80% consensus was determined as the host taxonomy. For the third approach, a TYMEFLIES MAG that contained the scaffold where the provirus was located was determined as the host.

If a viral genome was not assigned a host by any of the three approaches while it was placed in a species with the other member(s) assigned, the lowest common ancestor (LCA) of the host taxonomy for this species was used. Note that only the provirus, AMG host prediction, and “blast” or “CRISPR”-based iPHoP results were used to get host predictions from other species members. All the results were labeled with corresponding host taxonomy prediction approaches. The overlapped host taxonomies were resolved based on the following priority^3^: 1) provirus within a host genome; 2) “blast”-based iPHoP result; 3) “CRISPR”-based iPHoP result; 4) AMG match to host genome; 5) “iPHoP-RF” result; 6) derived from species host taxonomy.

### Auxiliary metabolic gene summary

The auxiliary metabolic genes (AMGs) identified by VIBRANT in the above section were first filtered according to the following criteria: (1) Edge-located AMGs (AMGs located at either end of a scaffold) were filtered. (2) AMGs that had any KEGG v-score or Pfam v-score (assigned by VIBRANT) ≥ 1 were filtered. (3) AMGs with flanking genes (four genes on either the upstream or downstream sites) having a KEGG v-score < 0.25 were filtered. (4) AMGs with annotation by COG category as “T” or “B” were filtered. The filtered AMGs were then summarized by adding the information, including date and season, KO hit and name, Pfam hit and name, and KEGG metabolism, pathway, and module. The AMG cluster occurrence (the number of metagenomes in which an AMG cluster can be found) and abundance (the mean normalized abundance of AMG cluster containing viruses in the metagenomes that this AMG cluster can be found) were obtained by summarizing AMG cluster containing viruses and were used to make scatter plots to find potential relationships (R library “ggpmisc”).

The summary of AMG cluster abundance for each season (a season from 20 years combined) and each year-season (a season from each year; e.g., “2000-Spring”) was conducted using the following steps: 1) Calculate AMG cluster abundance (normalized by 100M reads per metagenome) in each metagenome. 2) Calculate AMG cluster abundance for each season by adding up AMG cluster abundances from all metagenomes of this season (the AMG cluster abundance was normalized by the number of metagenomes in this season). 3) Calculate AMG cluster abundances for each year-season by adding up AMG cluster abundances from all metagenomes of this year-season (the AMG cluster abundance was normalized by the number of metagenomes in this year-season). 4) Generate AMG cluster abundance trend plots for each season (20 years combined) and each year-season (resulting in 20 facets for all 20 years) using R (R library “ggpubr”).

### AMG cluster variation

In non-singleton species, we investigated the AMG cluster variation by calculating the presence ratio of the AMG clusters across all the viral members within the species. To investigate how the viral genome completeness influences AMG cluster variation within a viral species, we selected several important AMG clusters for analysis. Viral genomes were categorized into five completeness levels: “75-100% complete”, “50-75% complete”, “25-50% complete”, “0-25% complete”, and “NA”. For the species with its species representative genome containing a specific AMG cluster, we calculated the AMG cluster containing ratio (as a percentage) by dividing the number of AMG cluster containing viral genomes by the total number of viral genomes within each completeness category. The results that represent the relation of virus completeness to the AMG cluster containing percentage for species’ members across all selected AMG clusters were plotted using bar plots in R (R library “ggplot2”).

The influence of species size (number of viral genomes in the species) on the AMG cluster variation was analyzed by dividing the combinations of AMG cluster and species into four quartiles according to the species size. The combination of AMG cluster and species was used for AMG cluster variation analysis because some species can contain multiple AMG clusters. The mean AMG cluster presence ratio for each AMG cluster from the AMG cluster and species combinations of the 1^st^ quartile (75-100%) of AMG cluster presence ratio category (the highest presence ratio) with the species size in the 4^th^ quartile (the largest species size) was calculated. It was then plotted against the AMG cluster count fraction (the percentage of occurrences of a single AMG cluster among all AMG clusters within a species) to illustrate the relationship. The presence tables of AMG clusters for each season (20 years combined) were obtained for individual AMG clusters. They were compared to the available metagenomes for each season. The percentage of AMG cluster containing metagenome number over the total metagenome number in each season was calculated for each high occurrence AMG cluster (distributed > 400 metagenomes).

### AMG cluster carrying virus and host diversity

To get the alpha diversity of viruses and their hosts for each AMG cluster, we used the family level taxonomy. Viruses with uninformative assignments (e.g., “Unclassified”, “NA;NA”, and “o ;f ”) were excluded. To evenly reflect alpha diversity, 100 viruses with informative family assignment and 25 viruses with informative host family assignment were randomly selected for each AMG cluster. Any AMG clusters that could not meet the required number of viruses were excluded. Alpha diversities (represented by Simpson indices) of viruses and viral hosts were obtained by R (R library “vegan”), and they were plotted against AMG cluster occurrence to find potential relationships (R library “ggpmisc”).

### AMG coverage ratio, viral genome abundance, and MAG abundance calculation

The mapping reference for viral abundance calculation was the collection of viral species representative genomes. These genomes were also the longest among species members. The AMG counterpart gene-located microbial scaffolds from all metagenomes were also included to avoid potential mis-mapping of microbial reads to viral AMGs. The mapping process was conducted by Bowtie 2 using all metagenomic reads with default settings. The resulting bam files were subjected to viral abundance and microdiversity analysis by MetaPop v0.0.60^69^ using the settings of “--id_min 93 --snp_scale both” (gene files from the above VIBRANT analysis were used in place of self-annotation by MetaPop; modifications were made to the gene files to adapt them to MetaPop requirements).

A custom script “cov_by_region.py” was used to parse the site-specific depth file (within “04.Depth_per_Pos” directory of MetaPop result from the above section) to get the AMG coverage and viral scaffold coverage (excluding all the AMG regions). We obtained the normalized viral genome coverage by first calculating the average of all its scaffold coverage values (excluding all the AMG regions) and then normalizing it by setting each metagenome read number as 100M. Note that all scaffold coverages from a viral genome should pass the cutoff of ≥ 0.01; otherwise, we assigned this viral genome as “absent”.

After summarizing viral genome coverages across all the metagenomes, we set a custom viral genome presence cutoff as follows: coverage ≥ 0.33 and breadth ≥ 50%. Then, based on these “present” viral genomes, we obtained the corresponding viral genome coverage (or referred to as “abundance”) and AMG coverage values across all the metagenomes, as well as the AMG coverage ratios (AMG coverage divided by its corresponding viral genome coverage). Using the same criteria for scaffold coverage cutoff and viral genome coverage and breadth cutoffs, we calculated the viral genome abundance for the other non-AMG-containing viruses.

TYMEFLIES species representative MAGs were used as the mapping reference for conducting metagenomic read mapping using Bowtie 2 with default settings. CoverM was used to calculate contig abundance using the settings of “--min-read-percent-identity 93 -m metabat”. The MAG was assigned as “present” in each metagenome with a breadth cutoff of 10%. The MAG abundance was calculated by computing the average contig abundance using the ratios of contig length to genome length. The MAG taxa abundance (at the family level) for each season was summarized using similar methods described above (refer to the second paragraph of the section “Auxiliary metabolic gene summary”).

### Virus composition pattern analysis

The abundance of viruses at the family level across all metagenomes was summarized. Subsequently, we generated Non-metric MultiDimensional Scaling (NMDS) plots by R (R library “vegan” and “ggplot2”). These plots represent the ordination of metagenomes based on pairwise distances of virus composition among all the metagenomes. According to the metagenome to season corresponding relationship, the ANOSIM test was conducted to inspect whether there was a statistical difference among metagenomes from different seasons using R (R library “vegan”) with options of “distance = ‘bray’, permutations = 9999”.

### Virus and host association analysis

For the calculation of Cyanobacteria virus and host abundances, we mainly focused on the three Cyanobacteria groups: *Cyanobiaceae*, *Microcystis*, and *Planktothrix*. Within each group, we computed the abundances of MAGs, *psbA*-containing viral genomes, and non-*psbA*-containing viral genomes, specifically from the 0 day of early summer for each year. Subsequently, we employed an interpolation function to generate abundance profiles for each year, with five-day intervals. To plot the mean curves for virus and host abundances, we initially obtained the mean values of abundance percentages (normalized by the highest abundance within one year) for each time point. We then multiplied these mean abundance percentages by the highest abundance within that respective year. This approach was implemented to mitigate the impact of substantial abundance fluctuations from year to year. The mean value for each time point was calculated based on valid abundances from at least three years. Subsequently, for each of the Cyanobacteria groups, we plotted the mean abundance curves for all viruses and hosts in a single line chart frame using Python 3 (Python library “matplotlib”).

For the calculation of methanotroph virus and host abundances, we mainly focused on the four methanotroph genera: *Methylocystis*, UBA6136, *Methylomonas*, and UBA10906. Similarly, within each genus, we computed the abundances of MAGs, *pmoC*-containing viral genomes, and non-*pmoC*-containing viral genomes, specifically from the 0 day of late summer for each year. The remaining methods were consistent with those described in the preceding paragraph to plot the mean abundance curves for all viruses and hosts. For the calculation of *Nanopelagicales* virus and host abundances, we mainly focused on two genera: *Planktophila* and *Nanopelagicus.* Similarly, within each genus, we computed the abundances of MAGs, *katG*-containing viral genomes, and non-*katG*-containing viral genomes, specifically from the 0 day of clearwater for each year. The remaining methods were consistent with those described in the preceding paragraph to plot the mean abundance curves for all viruses and hosts.

To calculate the viral-to-host abundance ratios for each group mentioned in the previous paragraphs, we considered pairs of virus and host abundances that met specific criteria. Specifically, at each time point, both the virus and host abundance percentages were required to be non-null (not “nan”) and greater than 10% to be considered valid pairs. Subsequently, box plots were created to show the viral-to-host abundance ratio distribution for each group using R (R library “ggplots2” and “scales”) with the ratio range displayed with log-transformed values.

### Microdiversity analysis

The microdiversity parameters were parsed based on the results of MetaPop (the “Microdiversity” folder). Only viral scaffolds that passed the requirement of microdiversity calculation (breadth ≥ 70% and depth ≥ 10) were taken into consideration. Similar to the methods described in the above sections, the following microdiversity parameters for each metagenome, each year-season, each season, and/or each year were calculated and summarized accordingly: nucleotide diversity (pi) for viral genome and viral genes, SNP density for viral genomes, rates of non-synonymous (pN) and synonymous (pS) polymorphism (pN/pS) for viral genes, and fixation index (*FST*) for viral scaffolds. The correlations between viral genome abundance with nucleotide diversity and SNP density were calculated by Python 3 using Spearman’s rank correlation test with the *p*-value provided.

To investigate the populational genetic alteration between summer and winter (Late Summer vs Ice-on) and between the beginning and ending years (2000-2003 vs 2016-2019), we conducted similar MetaPop analyses by aggregating the relevant metagenomic reads. Likewise, we calculated and summarized the four microdiversity parameters accordingly.

Yearly SNP allele frequency was calculated by parsing SNPs across the full length of the viral genome^6^. The “reference” alleles were chosen to be the predominant alleles of viral genomes in 2018. The choice was made because 2018 has the most metagenomes (*n* = 44) in the latter years and a yearly-changing trend from the beginning to the ending years can be depicted simply. SNP allele frequency was the percent of reads matching the “reference” allele at each SNP locus.

Yearly gene frequency was calculated to reflect the gene relative abundance change in the viral species population along the time-series. Gene frequency was estimated as the coverage of each gene divided by the mean coverage of all other genes in the genome^6^. To set the detection limit for genome coverage, the mean coverage of all genes in the genome was required to be ≥ 5.

Genes of length < 450 bp were excluded from the analysis. In addition, to avoid the coverage variation influenced by the “all-to-all” read mapping method (the default setting of Bowtie 2), the positions within the first and last 150 bp of a scaffold were excluded from coverage calculations. To get a statistically meaningful gene frequency, there was another requirement that the gene number in a genome with a valid coverage (not “NA”) should be over 50% of the total gene number. A gene frequency of 1.0 indicates that statistically compared to the other genes in the genome each virus in the population encodes one copy of the gene. Gene frequencies were considered significantly increased or decreased within a population if the change in gene frequency was ≥ 1.0. The yearly-changing trend of gene frequency was fitted to the linear regression by Python 3.

