## Supplementary Information for "A 20-year time-series of a freshwater lake reveals seasonal dynamics and environmental drivers of viral diversity, ecology, and evolution"

### Supplementary Methods

#### Fixation index calculations for microdiversity analysis

We introduced two groups of comparisons to investigate the *F_ST_* of each viral scaffold. First, all summer and winter metagenomic reads were combined (summer and winter metagenomes were selected when the season was assigned as “Late Summer” and “Ice-on”); second, metagenomic reads from the first four years (2000-2003) and the last four years (2016-2019) were combined. After read mapping, the resulting bam files were subjected to microdiversity analysis by MetaPop v0.0.60^1^ using the settings of “--id_min 93 --snp_scale both” (gene files from the VIBRANT analysis were used in place of self-annotation by MetaPop; modifications were made to the gene files to adapt them to MetaPop’s requirements).

The *F_ST_* (fixation index) values of each viral scaffold between metagenomes/samples were parsed from the results of MetaPop in the “Microdiversity” folder^1^. Viral scaffolds with *F_ST_* > 0.15 were identified as significantly different between groups. For viral genes located in these viral scaffolds, the following requirements were used to determine the selected viral genes: 1) Nucleotide diversity (pi) in the winter group > nucleotide diversity (pi) in the summer group or nucleotide diversity (pi) in 2000-2003 > nucleotide diversity (pi) in 2016-2019; 2) Gene N/S SNV ratio (pN/pS) in the winter group < gene N/S SNV ratio (pN/pS) in the summer group or gene N/S SNV ratio (pN/pS) in 2000-2003 < gene N/S SNV ratio (pN/pS) in 2016-2019.

#### RNA extraction and metatranscriptome processing

The water samples were collected approximately every 2 hrs from 6:00 am, August 20 to 6:00 am, August 21, 2015, for a diel cycle and used for RNA extraction and metatranscriptome processing. The sampling location was the same as described in the “Methods and Materials” section of the main text. The sampling depth was 0.5 m. These water samples were subjected to sample filtration, RNA extraction, rRNA depletion, addition of an internal standard, and sent to the University of Wisconsin – Madison Biotech Center for sequencing. The details of cDNA synthesis and library preparation are described at <https://github.com/McMahonLab/OMD-TOIL>. Specifically, metatranscriptomic reads from four samples (4 hrs past sunrise, 6 hrs past sunrise, 16 hrs past sunrise, and 20 hrs past sunrise) were combined and used for downstream analysis. The quality filtering and trimming were conducted by fastp v0.23.0^2^ using default settings, and *in silico* rRNA removal was conducted by SortMeRNA v4.3.6^3^ using default settings and rRNA database.

Four water samples from July to October 2020 were collected in the same location as described in the “Methods and Materials” section of the main text. The sampling depth was 5 or 10 m. The water was filtered in the lab 1-5 hrs after sampling through a 0.2 μm filter. Briefly, RNA extraction was performed according to a phenol-chloroform protocol described previously^4^. Specifically, each filter was placed into a bead-beating tube with 1 mL Trizol added and bead-beaten for 1 min at medium speed. The internal standard was added and mixed^5^. Each tube was centrifuged for 1 min. The resultant supernatant was collected, and 300 μL chloroform was added, followed by thorough mixing through inversion for 1 min. The mixture was then incubated at room temperature for 3 min. Subsequently, the mixture was subjected to centrifugation at 4°C with a speed of 13200×g for 15 mins. The supernatant was transferred to a new tube and mixed with cold 70% ethanol. The RNA extraction process was performed using the RNeasy Kit (Qiagen). The presence of residual DNA was eliminated using the TURBO DNA-free™ Kit (Invitrogen). The quantity of RNA was determined using the Qubit RNA Kit (Invitrogen). The RNA was subjected to SeqCenter (Pittsburgh, USA; previously MiGS) for sequencing using a standard RNA-seq platform and results processing routine. The resulting metatranscriptome datasets were subjected to *in silico* rRNA removal using sortmeRNA^3^.

#### Metatranscriptome mapping

For the 2015 diel cycle metatranscriptome datasets, the combined metatranscriptome with four samples collected during the diel cycle from August 20 to August 21, 2015 was used for downstream mapping. Two metatranscriptomic mapping approaches have been conducted. First, the mapping reference for the combined 2015 diel cycle metatranscriptome was the gene sequences from the two temporally closest Lake Mendota water sample metagenomes (IMG: 3300042395 collected on August 19, 2015 and IMG: 3300042470 collected on August 22, 2015). Secondly, the mapping reference for the combined 2015 diel cycle metatranscriptome was the collection of all viral species representative genomes with auxiliary metabolic genes (AMGs). In order to conduct competitive mapping, the AMG counterpart genes and their flanking regions from all Lake Mendota metagenomes were also added to the mapping reference.

For the metatranscriptomes of four water samples from July to October 2020, similarly, two metatranscriptomic mapping approaches were conducted. First, the mapping reference for each metatranscriptome was the gene sequences from the temporally closest (in month and date) Lake Mendota water sample metagenome (metagenome IMG: 3300044631 collected on July 24, 2018 for metatranscriptome ME_2020_07_24_5m_C; metagenome IMG: 3300044608 collected on August 6, 2018 for metatranscriptome ME_2020_08_05_10M_B; metagenome IMG: 3300042355 collected on August 25, 2018 for metatranscriptome ME_2020_08_25_10M_B; metagenome IMG: 3300034116 collected on October 24, 2018 for metatranscriptome ME_2020_10_19_5M_B). Secondly, similar to the approach described in the above paragraph, the mapping reference for each metatranscriptome was the collection of all viral species representative genomes with AMGs. The AMG counterpart genes and their flanking regions from all Lake Mendota metagenomes were also added to the mapping reference for competitive mapping.

Bowtie2 v2.4.5 was used for metatranscriptomic mapping with default settings. The resulting bam files were filtered with at least 50 bp aligned length and 0.97 read identity using a custom python script. The coverage of each contig was calculated by CoverM v0.6.1 with the option of “--methods rpkm”. The resulting coverage value for each contig represents the reads per kilobase of transcript per million reads mapped (RPKM). A viral genome was determined to have positive expression when ≥ 50% of its genes had positive expression.

### Supplementary Results

#### AMGs revealed an unprecedented comprehensive functional repertoire associated with freshwater viruses

To have a comprehensive view of viral auxiliary metabolisms, we summarized all viral AMGs with KEGG-based pathways/functions (Fig. S2). Beyond photosystem II domains, we also detected both photosystem I gene cassette components (*psaC*, *D*) (Fig. S2a)^6^. We detected *pmoC* for methane oxidation^7^, *frmA* for formaldehyde oxidation, and *FdH* and *fdhA* for formate oxidation; meanwhile, two formaldehyde assimilation pathways – serine and ribulose monophosphate pathways – were also discovered (Fig. S2b). To our current knowledge, these findings are novel for freshwater limnic environments ^7, 8, 9, 10^. High *glnA* occurrence and abundance indicated that viruses can boost the ammonia catabolism to generate precursors for pyrimidine synthesis (Fig. S2c). Viral AMGs participated in both the Wood–Ljungdahl pathway and reductive citrate cycle for CO_2_ fixation (Fig. S2d). In sulfur and related metabolism, high occurrence and abundance viral *cysC* and *cysH* participated in assimilatory sulfate reduction (Fig. S2e). Sulfur AMGs were widely present in pathways of viral-assisted cysteine synthesis/breakdown to sulfide, methionine salvage pathway, methionine degradation pathways, sulfonate conversion to sulfite, and thiosulfate reduction to sulfide (Fig. S2e). This suggested that the viral organic and inorganic sulfur auxiliary metabolism can produce sulfide as the end product. As a multifunctional agent, sulfide can participate in various aspects of host metabolisms and viral processes^11^, e.g., benefitting host survival/growth, amino acid synthesis or protein function, virion assembly, DNA/RNA modification, gene modification, enzyme function, etc. Thus, AMG-assisted/enhanced sulfide production can provide fitness benefits to viruses. The existence of viral *dmdC* suggested that viral infection can enhance the breakdown of dimethylsulfoniopropionate (DMSP). Viral *metK* gene for S-adenosyl methionine (abbreviated as AdoMet or SAM) production suggested that viruses can stimulate the generation of intermediaries for radical SAM enzymes, a group of enzymes utilized a [4Fe–4S] cluster and SAM to initiate a diverse set of radical reactions^12^.

All gene components of the Pentose Phosphate pathway were found (Table S6). The Pentose Phosphate pathway can generate precursors for the biosynthesis of nucleotides (through purine metabolism) and cofactors (e.g., riboflavin). It also generates NADP^+^/NADPH for biosynthesis and production of deoxyribonucleotides and fatty acids as a reductant^13^. Both the existence of viral AMGs in purine and pyrimidine metabolisms indicated that viruses boosted the synthesis of nucleotides for virion production^14, 15^ (Fig. S2f). Amino sugar and nucleotide sugar metabolism were enriched in phage-infected *Pseudomonas* virocells, which were postulated to link with the corresponding enriched viral AMGs^14, 16^. These metabolisms lead to the biosynthesis of peptidoglycans and lipopolysaccharides required for cell growth and division (Fig. S2g). The high occurrence and abundance of viral *ahbD* for heme synthesis indicated that viruses boosted the production of heme, assisting generation of heme peroxidase^17^ (Fig. S3i). As photosynthetic organisms fix CO_2_ under high-light conditions, they produce reactive oxygen species (ROS) such as H_2_O_2_ when electron transfer outpaces electron consumption^18^. The virus-assisted production of heme and heme-containing peroxidase/catalase (e.g., catalase-peroxidase encoded by *katG* and catalase encoded by *katE*, Fig. S2n) can help alleviate the host from damage by ROS. The biologically-generated ROS accumulation may reach its peak as the total photosynthetic biomass achieves a peak in warm, high-light irradiated epilimnion during summer^18^. The observed patterns that viral *ahbD* and *katG* abundances both peaked in clearwater and early summer (Dataset S1) supported this assumption. Recent culturing of the ubiquitous freshwater actinobacterial acI lineage depended on supplying catalase-peroxidase^19^. This suggested that previously yet-to-be cultured freshwater lineages were limited in self-production of catalase-peroxidases. By encoding *katG*, viruses can complement hosts to overcome ROS stresses^19^.

Virus-assisted heme generation can also promote the generation of cytochromes to replenish the redox cofactor pool for energy production (Fig. S2i)^13^. Cobalamin coenzyme participates in the tetrahydrofolate (THF)-mediated methyl-group transfer to acetyl-CoA in the Wood–Ljungdahl pathway for CO_2_ fixation. The high occurrence and abundance of *cob* genes indicated that virus-mediated metabolic redirection of hosts for boosting carbon assimilation (Fig. S2i). This can benefit viruses by synthesizing the required building blocks (e.g., nucleotides and proteins). Porphyrin and heme metabolism indicated viral-encoded AMGs (e.g., *bchE*, *ahbCD*, *HO*, etc.) can potentially enhance the biosynthesis of chlorophyll a and phycocyanobilin (a tetrapyrrole chromophore linked to phycobiliproteins, such as phycocyanin, for light harvesting^20^) (Fig. S2i), providing the building blocks of the photosynthetic machinery for hosts.

Virus-assisted folate biosynthesis pathway is involved with the synthesis of tetrahydrobiopterin (BH_4_), tetrahydrofolate (THF), and molybdopterin cofactor (Moco) (Fig. S2j). They are all important cofactors/prosthetic groups within enzymes for important reactions, e.g., the Wood–Ljungdahl pathway and serine pathway for formaldehyde assimilation (THF), production of nitric oxide from L-arginine (BH_4_), nitrate reduction, glyceraldehyde-3-phosphate dehydrogenation, carbon monoxide oxidation, formate oxidation, thiosulfate reduction etc. Viral-encoded AMGs were also associated with the biosynthesis of queuosine, which can improve the efficiency and accuracy of translation^21, 22^. Viruses encoded AMGs involving the nicotinate and nicotinamide metabolism, boosting the production of NAD^+^/NADH and NADP^+^/NADPH, two common coenzymes as redox intermediaries for various reactions in the cell^23^ (Fig. S2k). Riboflavin metabolism can generate flavin mononucleotide (FMN) and flavin adenine dinucleotide (FAD), which are important redox coenzymes participating in electron transport reactions^13^ (Fig. S2l). The products from the Pentose Phosphate Pathway (Ribulose-5P) and purine metabolism (GTP) can supply the precursors for coenzyme biosynthesis in riboflavin metabolism (Fig. S2g).

Beyond all the metabolisms/pathways, there were still other AMG clusters with high occurrence and abundance (Fig. S2m). Many participated in cell wall component biosynthesis (nucleotide sugar, lipopolysaccharide, peptidoglycan, and arabinogalactan), coenzyme biosynthesis (biotin and ubiquinone), and ROS stress alleviation (glutathione peroxidase^24^) as shown above. Additionally, many other AMG clusters were involved in energy generation (Fig. S2n), e.g., the TCA cycle and oxidative phosphorylation. Viruses can boost hosts in various carbon uptake processes, including alcohol, aromatics, sialic acid, galactose, pectin, starch/glycogen, cellulose, and mannan (Fig. S2n). Galactose, pectin, starch, cellulose, and mannan are important components of plant- or phytoplankton-derived polymeric or oligomeric carbohydrates^25^. The carbohydrates released by these photosynthetic organisms can contribute substantial amounts to the dissolved organic matter in freshwater lakes^25^. Together, these findings indicate that viral-encoded auxiliary metabolisms can fuel the utilization of carbohydrates from primary production (originating from plants and phytoplankton) to generate both energy and material for virion production, similar to previous discoveries of viral AMGs in soil^26, 27^.

#### AMG cluster variation in viral species

We examined the variance of AMG clusters in members of a viral species to understand their distribution across viral populations better. AMG cluster variation (presence ratio among all the members) in all species revealed that ~30% of the AMG cluster and species combinations (sometimes, multiple AMG clusters were present in a given species) had high AMG cluster presence ratios (distributed in the top 75-100% quartile) (Fig. S3a). The AMG cluster variation pattern remained generally consistent regardless of species size (along the x-axis; Fig. S3a).

We then examined the AMG cluster and species combinations from the 1^st^ quartile (75-100%) of the AMG cluster presence ratio category (the highest presence ratio) with species size in the 4^th^ quartile (the largest species size) to find out the distribution patterns for different AMG clusters (Fig. S3a). Across all species, high occurrence AMG clusters had higher AMG cluster presence ratios (mostly > 95%) and abundance (with AMG cluster count fraction mostly > 1%). This indicates that high occurrence AMG clusters tended to be consistently carried by multiple viruses across samples. Furthermore, when 20 years of time-series samples were aggregated, they were distributed across different seasons with high presence ratios within metagenomes (Fig. S3b). Overall, this indicates that high occurrence AMG clusters were distributed widely regardless of which viruses carried them and seasonality changes (and potentially the underlying environmental factors).

#### Temporally variable viruses have a high contribution to the AMG pool

To gain a better ecological and evolutionary understanding of viral populations with biogeochemical significance, we used the 20-year time-series metagenomic dataset to conduct coverage analyses. Consistent with the seasonal distribution patterns of AMG cluster abundance (Dataset S1), the total abundance of *psbA*-containing viral species peaked in late summer (Table S7). To conduct comparative analyses, we then chose late summer as the representative for each year. Time-series results showed that four viral species had a high yearly occurrence (≥ 15 out of 20) (Fig. S5a) and were persistent species throughout the 20 years. However, the membership of high AMG abundance species was observed to be inconsistent with high occurrence species (Fig. S5a). Two of the above four species had high AMG abundance (relative abundance ≥ 10%) (Fig. S5a). Similarly, *pmoC-*, *katG*-, and *ahbD*-containing viral species with high occurrence also differed from the high AMG abundance species (except for one *katG*-containing viral species and two *ahbD*-containing viral species) (Fig. S5b, S5c, S5d). Collectively, this suggests that persistent AMG-containing viral species are not typically the most abundant each year; instead, the pool of AMG abundance is largely influenced by viral species that vary annually. Since AMGs contribute to biogeochemical functions by regulating host activity, this finding suggests that temporally variable viral species may have a significant impact on crucial biogeochemical processes, such as photosynthesis and methane oxidation. Their abundances might fluctuate due to annual variations in environmental and host conditions. Further investigations are encouraged to unveil the underlying mechanisms governing these different abundance patterns.

#### Fixed viral genomes and selected genes

The Fixation index (*F_ST_*) measures the degree of population differentiation between two populations. To examine genetic variations in populations across temporal and seasonal shifts, we utilized MetaPop to identify viral scaffolds with *F_ST_* > 0.15^1, 28^, which indicates substantial differentiation, between the time intervals of 2000-2003 and 2016-2019, as well as between summer and winter seasons. The collection of all viral species representatives containing AMGs was used as the mapping reference for the populational analysis. In total, we identified 255 and 802 viral scaffolds with *F_ST_* > 0.15 for the comparisons of the time intervals of 2000-2003 and 2016-2019 and the summer and winter seasons.

We proceeded to analyze the microdiversity patterns of individual viral scaffolds in search of selected genes across temporal and seasonal shifts. Genes displaying lower nucleotide diversity (pi) and higher pN/pS values in 2016-2019 or summer samples compared to 2000-2003 or winter samples were identified as selected genes. These genes experienced positive selection during temporal and seasonal shifts, thus, leading to reduced nucleotide polymorphism and increased positive selection strength. Additionally, we filtered these viral scaffolds with *F_ST_* > 0.15, applying an extra criterion that more than 10% of the viral genes should be identified as selected genes. This process yielded 19 and 80 viral scaffolds for the comparisons between the time intervals of 2000-2003 and 2016-2019, as well as the summer and winter seasons, respectively. These represent the final set of fixed viral scaffolds and selected genes across temporal and seasonal shifts (Table S10).

When scrutinizing the functional characteristics of these selected genes on the fixed viral scaffolds, we found that 60-70% of them were assigned to the unknown functional category. This was followed by genes assigned into DNA/RNA replication/modification (13-18%), auxiliary metabolism (6-7%), and viral hallmark/structural (3-9%) functional categories. For the DNA/RNA replication/modification functional category, genes encoding important functions were selected, for instance, DNA polymerase, DNA primase, nuclease, AAA protein, DNA replication factor, DNA helicase, NTPase, ribonuclease, transcription initiation factor, and DNA synthesizing protein for DNA/RNA replication, and DNA methyltransferase, DNA recombination protein, DNA damage response protein, and DNA repair protein for DNA modification. For the auxiliary metabolism functional category, the selected gene functions were related to oxidative phosphorylation, linoleic acid metabolism, nucleotide metabolism, folate biosynthesis, porphyrin, heme, and cobalamin metabolism, amino sugar and nucleotide sugar metabolism, biosynthesis of nucleotide sugars, lipopolysaccharide biosynthesis, and biosynthesis of unsaturated fatty acids.

The DNA/RNA replication enzymes play a crucial role in the evolutionary significance of synthesizing viral nucleic acids and facilitating viral propagation. DNA modification enzymes, on the other hand, can lead to nucleotide changes, recombination, and epigenetic modifications. These processes are key contributors to viral genomic diversity and are pivotal drivers of viral evolution. DNA modification can have several important consequences. It can facilitate the adaptability of viruses to changing environments, such as, signaling a transition from a latent state to a lytic state^29^. Additionally, DNA modification can aid viruses in evading host defense systems, for example, the restriction-modification (RM) system, enhancing their ability to persist and propagate within host organisms^30^. The selected genes associated with auxiliary metabolisms encompass a wide range of functions, including energy conservation, nucleotide metabolism, cofactor biosynthesis, as well as amino sugar, nucleotide sugar, lipopolysaccharide, and unsaturated fatty acids metabolisms. These selected functions play vital roles in furnishing viruses with the necessary energy and substrates for replication, supporting diverse cellular metabolisms within host cells, and supplying essential components for cell wall biosynthesis and cell division. In summary, this reflects the pattern of genetic alterations within the viral population across temporal and seasonal shifts as driven by the process of viral fitness selection.

#### Viral genome and AMG expression

To investigate the viral genome and AMG gene expressions, we used five metatranscriptomic datasets that were obtained from water samples collected in either August 2015 or July to October 2020. We used two mapping references to examine the expression of viral genomes – (1) the corresponding metagenome(s) that are from samples collected on the same date of a year (or on the closest date of a year), (2) all virus species representative genomes. This approach can help investigate more viral genomes that were potentially captured in the metatranscriptomes. The viral genome and AMG expression showed that in positively expressed viral genomes, many AMGs were also detected with positive expressions (Table 1). This indicates that AMGs actively participate in the auxiliary metabolic functions within the host, suggesting that viruses actively contribute to the biogeochemical activities across the time-series.

**Table 1. The viral genome and AMG expression results.** Five sets of metatranscriptomic reads were mapped onto the gene collections of either the corresponding metagenome(s) or the all virus species representative genomes. The “diel_cycle_metaT” metatranscriptome dataset was combined from four water samples collected during the diel cycle from Aug 20 to Aug 21, 2015. The mapping reference labeled as “24hrs_metaT_ref” was made by combining the two metagenome datasets (3300042395 sampled on 8/19/2015 and 3300042470 sampled on 8/22/2015). A viral genome was determined to have a positive expression when ≥ 50% of its genes have positive expression.

| **Metatranscriptome mapping onto metagenome** | **No. of viral genomes with positive expression^a^** | **No. of viral genomes having positive AMG gene expression** |
| --- | --- | --- |
| diel_cycle_metaT___24hrs_metaT_ref | 1954 | 189 |
| diel_cycle_metaT___all_phage_species_rep^b^ | 7 | 7 |
| ME_2020_07_24_5m_C___3300044631 | 192 | 15 |
| ME_2020_08_05_10M_B___3300044608 | 5 | 5 |
| ME_2020_08_25_10M_B___3300042355 | 69 | 3 |
| ME_2020_10_19_5M_B___3300034116 | 106 | 9 |
| ME_2020_07_24_5m_C___all_phage_species_rep | 5 | 5 |
| ME_2020_08_05_10M_B___all_phage_species_rep | 4 | 4 |
| ME_2020_08_25_10M_B___all_phage_species_rep | 2 | 2 |
| ME_2020_10_19_5M_B___all_phage_species_rep | 7 | 7 |

^a^ Only the viral genomes containing at least one AMG were considered.

^b^ We only counted the viral genomes that contained four AMGs with biogeochemical significance (*psbA*, *pmoC*, *katG*, and *ahbD*) and have high occurrence and/or AMG abundance across 20 years.

### Supplementary Figure Captions

**Supplementary Figure S1. Viral scaffolds and genomes summary. a** Binned/unbinned scaffold percentage after binning by vRhyme and bin member number frequency for all bins (vMAGs). Bin member numbers only with frequencies > 1% were shown in the bar plot. Numbers of scaffolds and numbers of bins were labeled accordingly. **b** Length and completeness change after binning and CheckV quality to viral genome length distribution. Viral scaffold or/and vMAG (viral genome) numbers were labeled accordingly. “Viral scaffolds”: total viral scaffolds before binning; “vMAGs+unbinned scaffolds”: vMAGs and unbinned scaffolds after binning; “vMAGs”: vMAGs after binning; “Binned scaffolds (within vMAGs)”: binned scaffolds (the scaffolds that are in the vMAGs) after binning; “Unbinned scaffolds”: unbinned scaffolds after binning. The *p*-value indicates significance between comparisons. **c** The number of viral scaffolds, vMAGs, species, and genera. **d** Rarefaction curve of species-level vOTU numbers. Ten replicates with a random starting sample were made to generate error bars.

**Supplementary Figure S2. AMG metabolisms and functions. a** AMGs involved in photosynthesis, **b** AMGs involved in methane and related metabolism, **c** AMGs involved in nitrogen metabolism, **d** AMGs involved in CO_2_ fixation, **e** AMGs involved in sulfur and related metabolism, **f** AMGs in nucleotide metabolism, **g** AMGs in amino sugar and nucleotide sugar metabolism, **h** AMGs in pyruvate metabolism, **i** AMGs in porphyrin, heme, and cobalamin metabolism, **j** AMGs in folate biosynthesis, **k** AMGs in nicotinate and nicotinamide metabolism, **l** AMGs in riboflavin metabolism, **m** AMGs assigned to AMG clusters with other important functions (distributed > 300 metagenomes), **n** AMGs assigned to clusters with other important functions. The gene symbol and the corresponding enzyme name and CAZy ID for genes in **n** were depicted in brown together with the occurrence and abundance values (labeled as “occurrence|abundance”; occurrence, the number of metagenomes in which an AMG cluster can be found; abundance, the mean normalized abundance of AMG carrying viruses in the metagenomes in which this AMG can be found). Dotted arrows indicate steps that are not encoded by AMGs. Detailed information on each AMG cluster can be found in Supplementary Table S6.

**Supplementary Figure S3. AMG cluster variation in species and high occurrence AMG cluster distribution across different seasons. a** AMG cluster variation in species. The left bar plot represents the AMG cluster presence ratio pattern among all AMG cluster and species combinations. The x-axis indicates the size category of species and the number of AMG clusters and species combinations. The y-axis indicated the fractions of four quartiles of AMG cluster presence ratios. The right scatter plot represents the AMG cluster count fraction (the percentage of one AMG cluster being encountered among all AMG clusters within a species) to the mean AMG cluster presence ratio (the percentage that one AMG cluster appears among all members within a species) across all species. This scatter plot used the AMG cluster and species combinations of the 1^st^ quartile (75-100%) of the AMG cluster presence ratio category (the highest presence ratio) with the species size in the 4^th^ quartile (the largest species size), which was shown as the connection by dash lines. High occurrence AMG clusters (distributed > 400 metagenomes) were colored red, and other AMG clusters were colored green. **b** Seasonal distribution of high occurrence AMG clusters distributed > 400 metagenomes) across metagenomes. The percentage indicates the AMG cluster containing metagenome number over the total metagenome number in each season.

**Supplementary Figure S4.** Taxonomic distribution (classified into the family level) of predicted hosts for eight AMG clusters with low Simpson indices. Unclassified hosts were not depicted and low abundance families (with abundance < 5% in all eight AMG clusters) were integrated into a taxon named “The rest”.

**Supplementary Figure S5. Species and AMG abundance across the time-series.** Seasonal abundance distribution, species and AMG abundance percentage, and total species and AMG abundance across 20 years for *psbA*- (**a**), *pmoC*- (**b**), *katG*- (**c**), *ahbD*-containing (**d**) viruses were summarized. In each subpanel, high occurrence species were picked according to the occurrence across 20 years, high abundance AMGs were picked according to the non-zero mean relative abundance across 20 years, and the abundance for each year was represented by the season with the highest/second to the highest species abundance in each year (Late Summer for *psbA*, Fall for *pmoC*, Late Summer for *katG*, and Early Summer for *ahbD*). Species and AMG were colored in blue and orange, respectively. Star-labeled AMGs indicated the overlap of the high occurrence species and high abundance AMG in subpanels **a**, **c**, **d**. The abundance values (for both species and AMG) were normalized by 100M reads/metagenome. For *psbA*- and *ahbD*-containing viruses, only species with ≥ 20 occurrences out of 471 metagenomes were included in the analysis; for *pmoC*- and *katG*-containing viruses, only species with ≥ 5 occurrences out of 471 metagenomes were included in the analysis. The species and AMG abundance percentage calculation was based on the total occurrence-filtered viral species.

**Supplementary Figure S6. Phylogenetic tree of viral and microbial PmoA.** This PmoA tree was re-rooted by the bacterial AmoA (and AmoA-like) sequences. Gammaproteobacterial and Alphaproteobacterial PmoA groups were labeled with color boxes. The PmoA sequences originating from MAGs and viral genomes in this study were labeled yellow and blue, respectively. Additionally, MAG taxonomies were labeled in the sequences. The nodes with ultrafast bootstrap (UFBoot) support values ≥ 90% were labeled with black dots accordingly.
