## Supplementary figures and images for "A 20-year time-series of a freshwater lake reveals seasonal dynamics and environmental drivers of viral diversity, ecology, and evolution"

### Figure S1

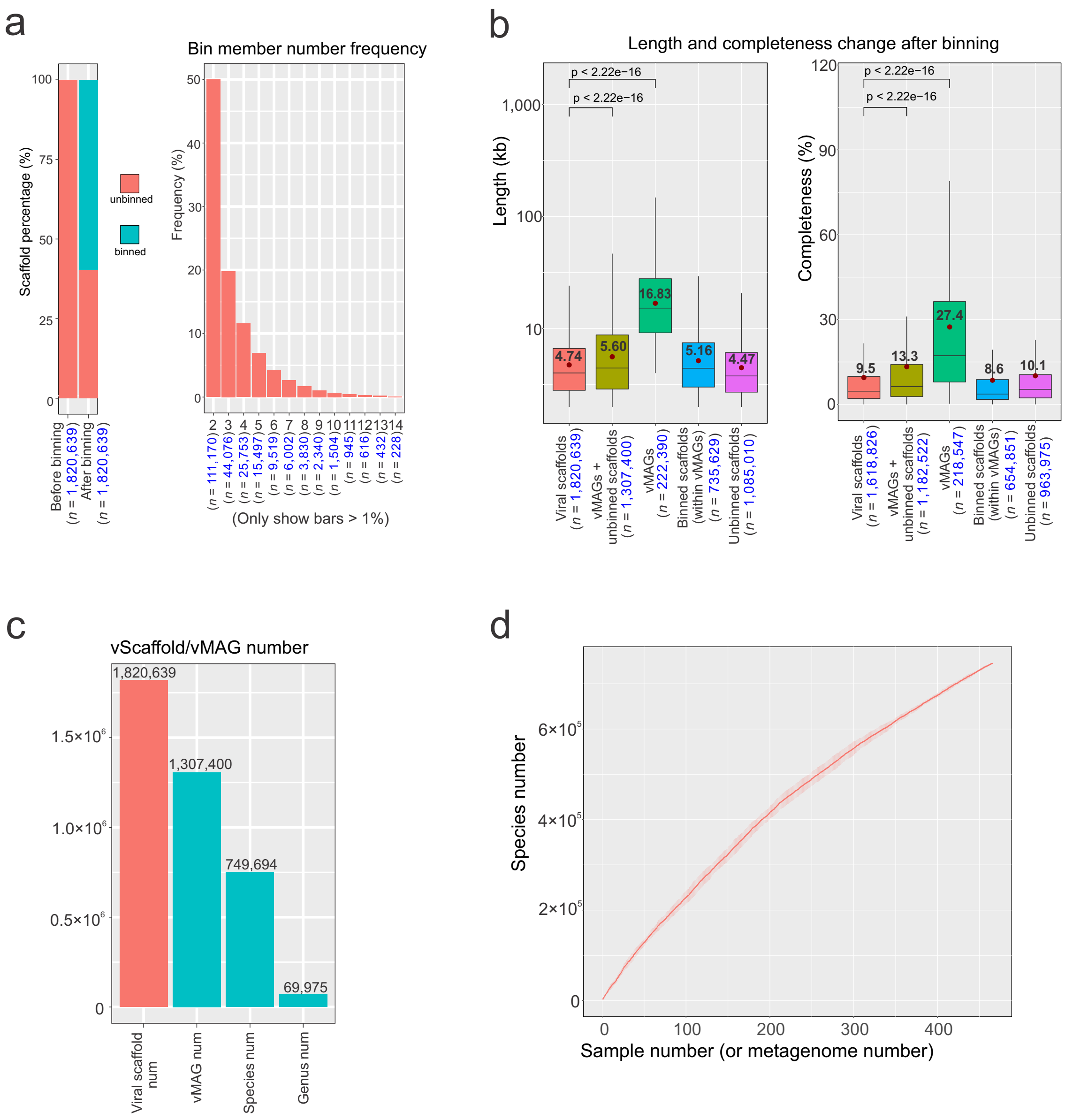

### Figure S2

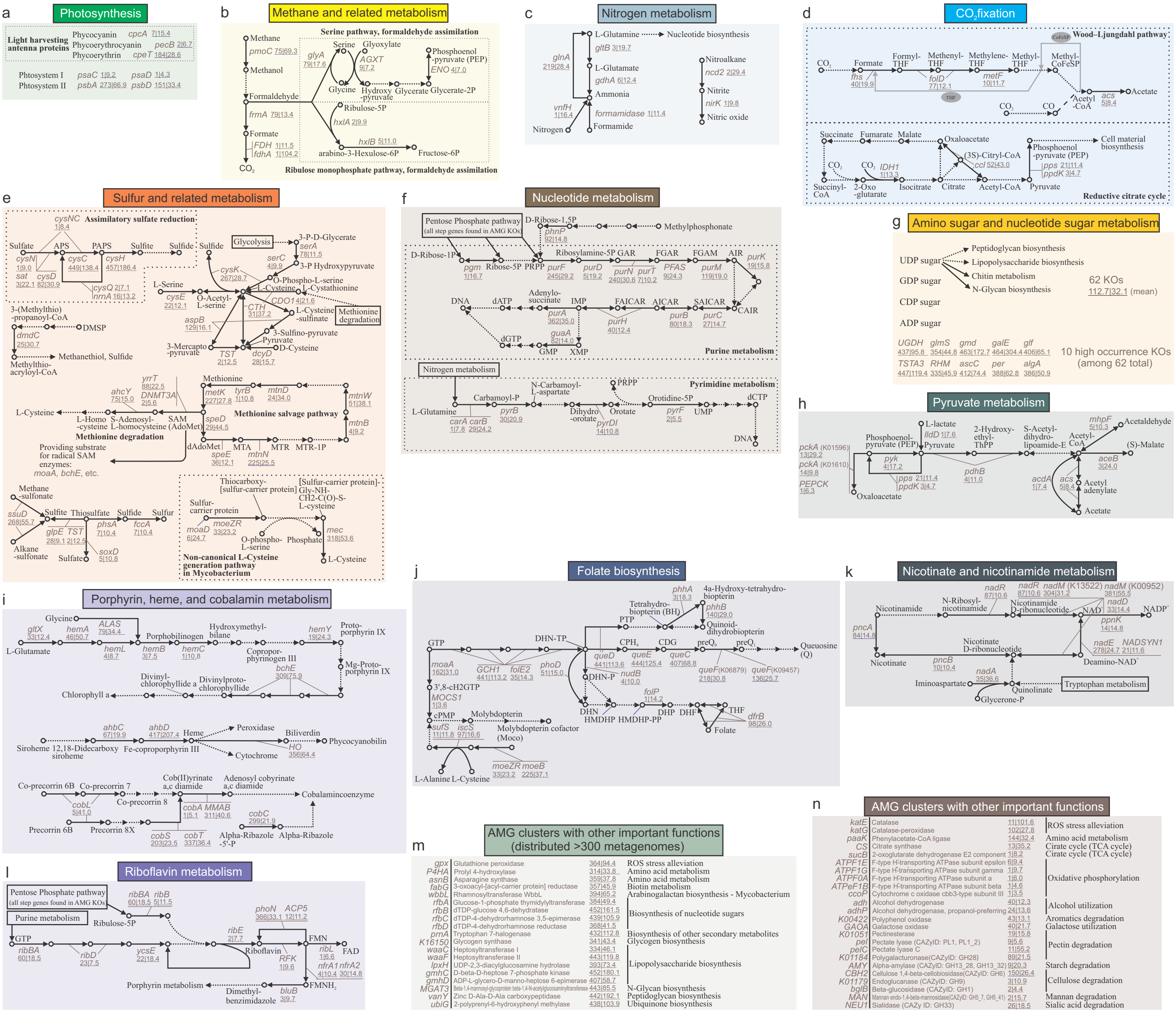

### Figure S3

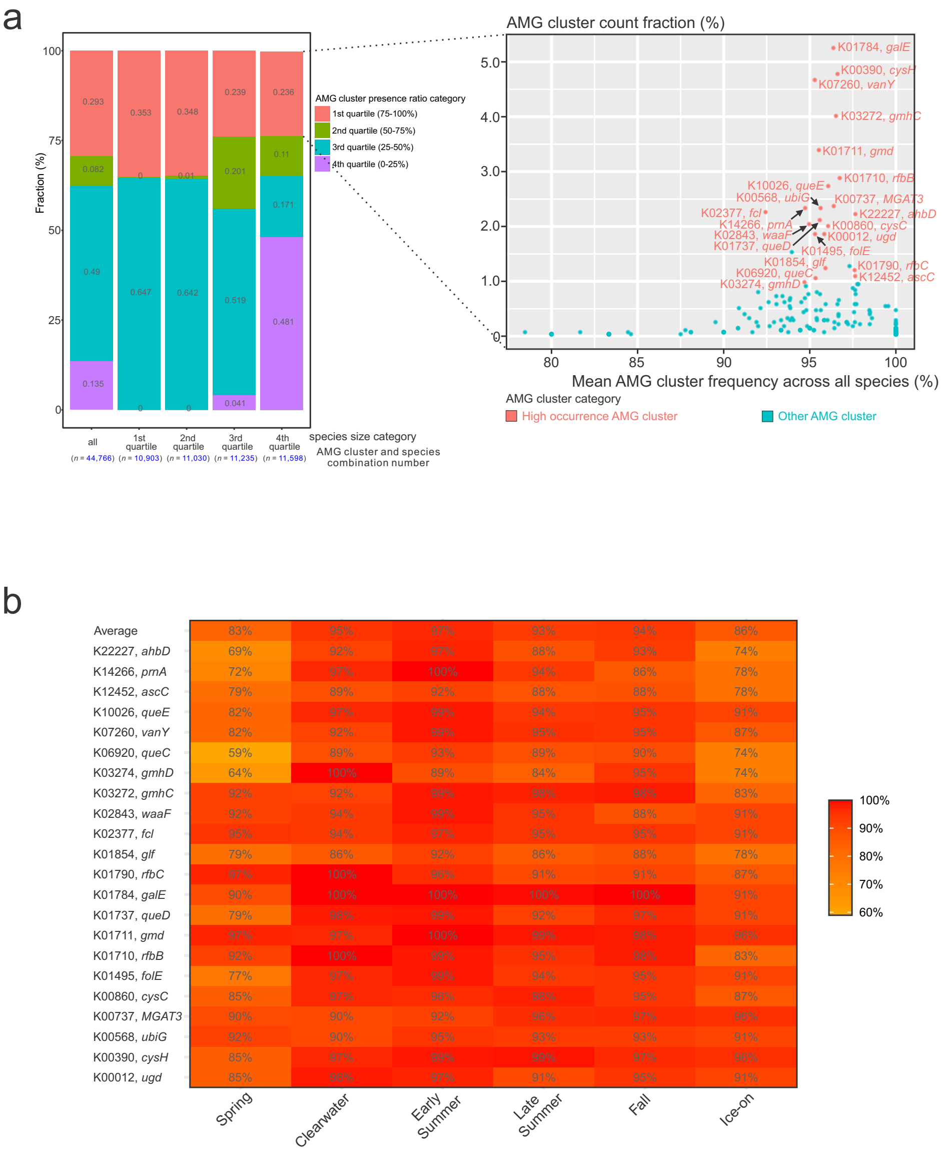

### Figure S4

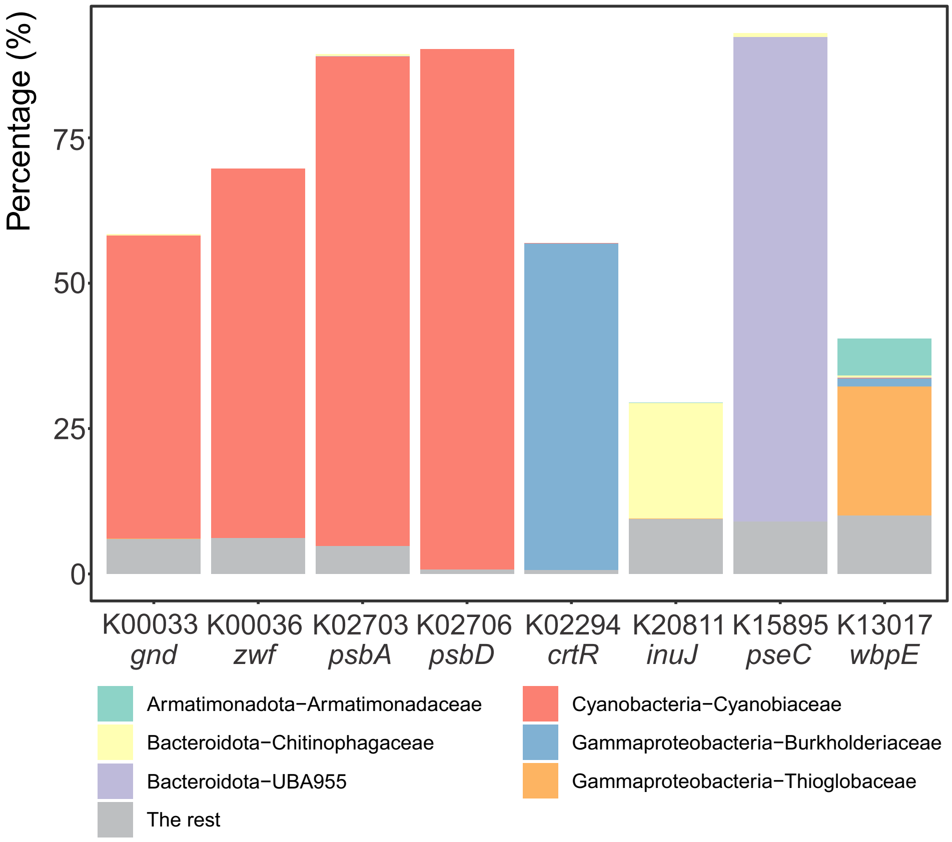

### Figure S5

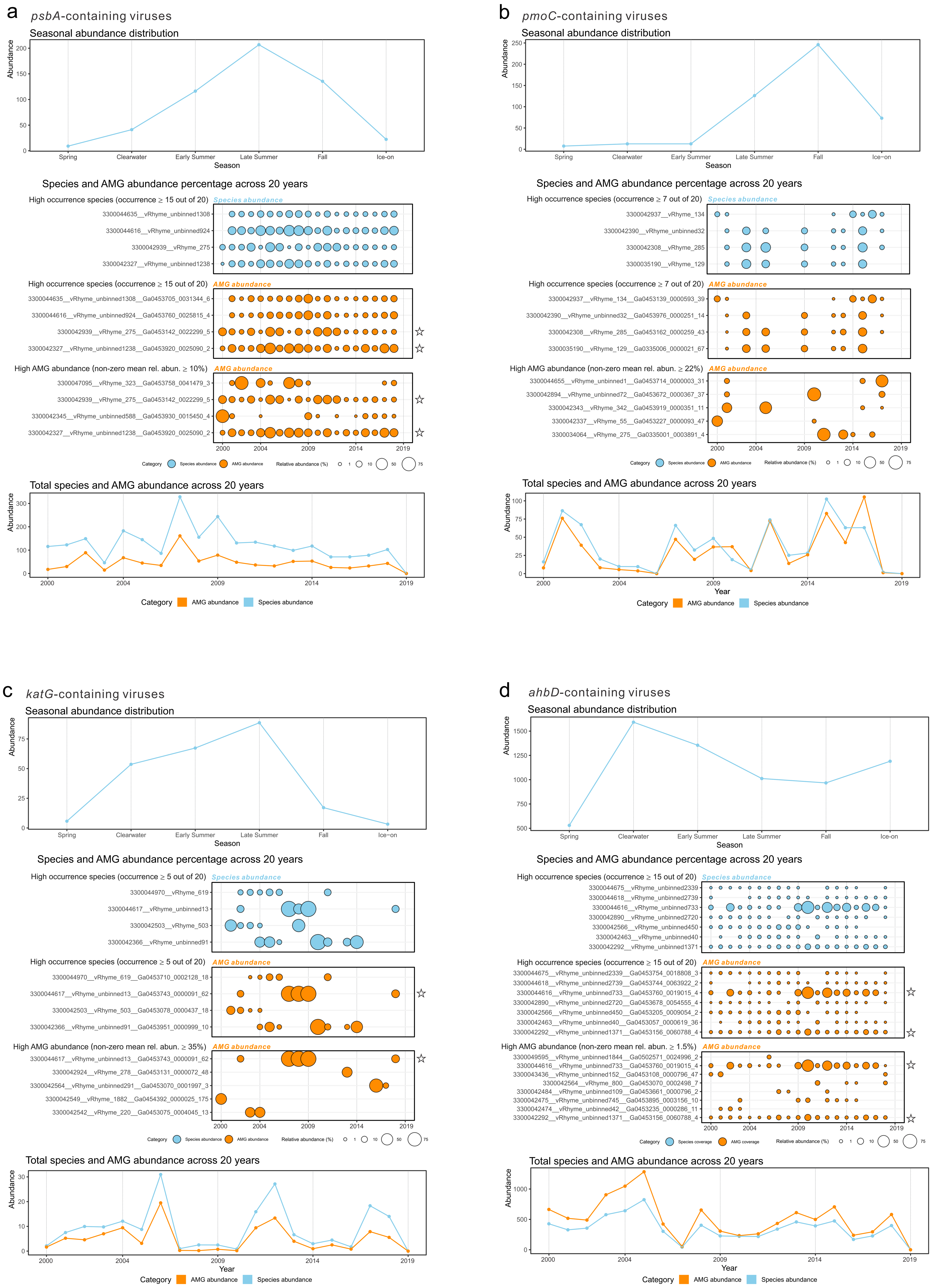

### Figure S6

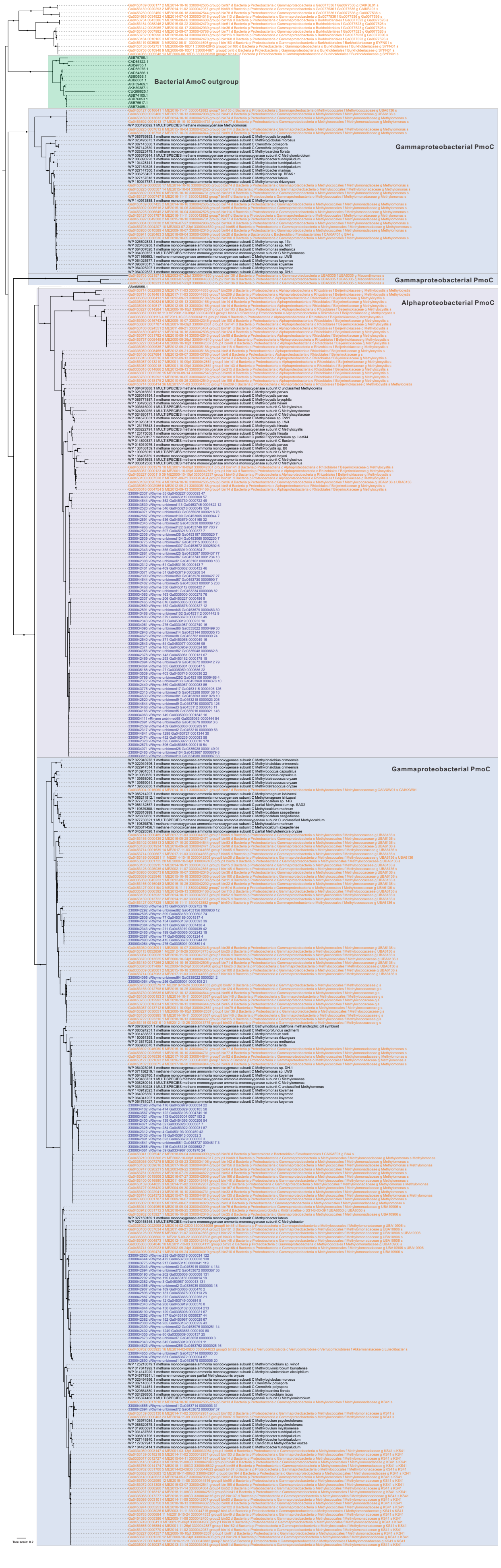
